# Dually-decorated palmitate-containing lipid nanoparticles for the targeted delivery of siRNAs against HER2 and Hsp27 in HER2+ breast cancer

**DOI:** 10.1101/2025.07.27.667055

**Authors:** Pedro Medina, Modesto Orozco, Montserrat Terrazas

## Abstract

Ionizable lipid nanoparticles (LNPs) have enabled significant advances in oligonucleotide therapeutics, culminating in the clinical approval of an siRNA-based formulation. Unfortunately, practical use of siRNA containing LNPs is still hampered by difficulties, one of them directing LNPs toward specific cells. To improve cell-specific delivery, we have developed a new generation of palmitate-containing ionizable LNPs functionalized with two distinct ligands: i) a targeting peptide (octreotide, Oct), for selective recognition of somatostatin receptor 2 (SSTR2)-expressing tumor cells, and ii) a cell penetrating peptide (penetratin, RK16) to enhance intracellular delivery. This dually-decorated LNP was used to co-deliver two siRNAs targeting key mediators of drug resistance in HER2+ breast cancer –Hsp27 and HER2– demonstrating strong potential for combinatorial RNAi-based therapies. The Oct moiety conferred high selectivity toward malignant SSTR2-overexpresing (SSTR2OE) HER2+ BC cells, even in heterogeneous environments containing non-tumor epithelial cells. Simultaneously, the RK16 peptide facilitated enhanced intracellular delivery, resulting in improved antiproliferative activity and a notable reduction in the SSTR2-overexpressing cell population compared to non-functionalized LNPs. Taken together, the structural design and biological performance of this novel class of LNP formulation establish it as a versatile and innovative delivery platform for ON-based targeted therapies. By simply altering the targeting peptide moiety and siRNA combination, this dually-functionalized LNP system could be readily adapted for a broad range of tumors and combination treatment strategies. Finally, the inclusion of palmitic acid in the LNP composition selectively contributes to cytotoxicity in palmitate sensitive cell lines, such as SK-BR-3, offering an additional therapeutic advantage when targeting these tumor subtypes.

## 1. INTRODUCTION

Oligonucleotide (ON) therapeutics,^1–9^ including antisense oligonucleotides (ASOs),^6^ short interfering RNAs (siRNAs),^7^ aptamers,^9^ and CRISPR-Cas systems,^8^ have revolutionized modern medicine. These therapies allow for the precise modulation or inhibition of gene expression across the entire human genome, making it possible to target proteins that are usually unreachable to small molecule drugs. By directly interacting with mRNA or genomic DNA, ON therapeutics provide promising treatment options for various pathologies, such as genetic disorders, viral infections, and cancers, with remarkable specificity and efficiency.

To overcome the major challenges in the clinical application of ON therapeutics –namely, poor biostability and limited cellular uptake– various chemical modifications and delivery platforms have been developed.^1–5^ These include sugar modifications such as 2’-*O*-Me,^10^ 2’-*O*-MOE,^11^ and 2’-F,^12^ as well as backbone alterations like phosphorothioate (PS) substitutions^13^ to improve nuclease resistance. Delivery strategies often involve conjugating the ON to specific ligands,^14–19^ such as *N*-acetylgalactosamine (GalNAc), which targets the asialoglycoprotein receptor^20^ on hepatocytes,^15,20^ or encapsulating the ON in lipid nanoparticles (LNPs).^4,21–24^ These LNPs typically consist of a polyethylene glycol (PEG)-lipid to prevent aggregation; cholesterol and a phospholipid [e.g., 1,2-distearoyl-*sn*-glycero-3-phosphocholine (DSPC) and 1,2-dioleoyl-*sn*-glycero-3-phosphoethanolamine (DOPE)] for structural integrity; and an ionizable cationic lipid (e.g., DLin-MC3-DMA; p*K*_a_ = 6.4)^25^ that facilitates ON encapsulation at acidic pH and endosomal escape upon cell entry.^23^ The choice of PEG-lipid (e.g., with C16 and C18 chains) also affects circulation times and biodistribution.^26^

These design strategies have led to the approval of twenty-two ON drugs to date,^2,5^ including GalNAc-conjugated siRNAs, a LNP-formulated siRNA, ASOs, aptamers, and a CRISPR-based agent –most of which are delivered to the liver, eye or cerebrospinal fluid.^2,5^

While the liver remains a primary target due to high blood flow and natural LNP uptake via ApoE-LDL receptor interactions,^27^ efforts to extend delivery beyond hepatic tissue are ongoing.^23,24,28,29^ With this aim, LNPs have been engineered with targeting ligands (e.g., antibodies, peptides, small molecules) to achieve selective delivery to organs such as the spleen, lung, pancreas and prostate.^23,29,30^ Examples include antibody-decorated LNPs (CD3, CD4 or CD163 antibodies) for spleen delivery,^31^ GALA peptide-modified LNPs for lung delivery,^32^ and Lyp-1^33^ or Glu-urea-Lys^34^ peptide-modified LNPs for tumor targeting. Additionally, a new class of targeting LNP platform – namely, selective organ targeting (SORT) LNPs; enhanced with tissue-specific ionizable lipids– is being tested for improved delivery to the lung, spleen and kidneys.^2,27,28,35–39^ Extending these studies to target different organs and even specific cell types inside the organ is one of the main challenges in the therapeutic nucleic acids field.

This is particularly critical in the context of cancer, where therapeutic resistance and tumor heterogeneity further complicate treatment. Beyond the challenges of administration, cancer’s inherent complexity –driven by its multiple subtypes^40^ and dynamic signaling networks^41^– often necessitates combination treatments to overcome resistance mechanisms.^41,42^ For instance, in HER2+ BC, where Hsp27 is frequently overexpressed and linked to poor prognosis and drug resistance,^43–46^ co-treatment with a small molecule Hsp27 inhibitor and trastuzumab –a monoclonal antibody that disrupts HER2 signaling by preventing HER2 homodimerization and promoting receptor internalization^47^– has been shown to restore trastuzumab activity in trastuzumab-resistant HER2+ breast cancer cells.^46^

In this context, herein we present a novel class of ON delivery systems designed for selective tumor recognition and enhanced intracellular uptake. Our approach employs ionizable LNPs functionalized with two distinct ligands: i) a targeting peptide, octreotide^48^ (Oct) –a cyclooctapeptide agonist of the endocrine hormone somatostatin that binds with high affinity to somatostatin subtype-2 receptor (SSTR2), which is highly expressed in many tumor cells^49^– and ii) a cell penetrating peptide, penetratin (RK16),^50,51^ which facilitates endosomal escape and intracellular delivery. Both peptides are conjugated to the PEG-lipid component of the LNP, enabling a stepwise mechanism of tumor targeting followed by efficient cellular entry.

Dually-functionalized LNPs were successfully employed to co-deliver siRNAs targeting Hsp27 and HER2 in HER2+ BC cells, resulting in potent anti-proliferative effects without cytotoxicity to non-tumoral cells. While several previous studies have reported the co-delivery of ONs tackling different molecular pathways –for example, T7 peptide-singly-modified LNPs co-administering anti-Bcl-2 and anti-Akt-1 ASOs demonstrated synergistic effects in A549 and KB cancer cells, as well as and in A549 xenograft models^52^– our system, to the best of our knowledge, represents the first example of a dually-functionalized LNP capable of co-delivering therapeutic ONs with superior selectivity and efficacy compared to LNPs featuring a single ligand.

## 2. RESULTS AND DISCUSSION

### 2.1. Design and synthesis of decorated LNPs

Our dually-modified LNP design incorporates the following components: i) a neutral lipid (DSPC; 1,2-distearoyl-*sn*-glycero-3-phosphocholine); ii) the ionizable lipid DLin-MC3-DMA; iii) cholesterol; iv) DPG-PEG (1,2-dipalmitoyl-*rac*-glycero-3-methyl polyoxyethylene-2000), a PEG-lipid with two 16-carbon chains; iii) Oct-modified DSPE-PEG_2000_ (Oct-DSPE-PEG; where DSPE-PEG is a PEG-lipid with two longer C_18_ alkyl chains favoring stable conjugate insertion); and iv) RK16-modified DSPE-PEG_2000_ (RK16-DSPE-PEG) (Fig. 1A,B). This LNP formulation will be used to encapsulate a 1:1 combination of Hsp27 and HER2 siRNAs, resulting in the dually-loaded, dually-functionalized formulation termed 2s-Oct-RK16-LNP (Fig. 1A). To support cellular uptake studies and provide experimental controls, we will prepare the following dually (Oct- and RK16)-functionalized LNP variants (Fig. 1A): i) C-s-Oct-RK16-LNPs, loaded with a single siRNA (a Cy5-labeled version of HER2 siRNA; Cy5-HER2 siRNA; red emission); ii) e-Oct-RK16-LNPs, empty LNPs; and iii) scr-Oct-RK16-LNPs, loaded with a scrambled (non-targeting) siRNA. To dissect the individual contribution of each ligand, singly-functionalized LNPs decorated with either RK16 or Oct will be prepared, in both empty and siRNA-loaded forms. These include: i) 2s-RK16-LNPs and 2s-Oct-LNPs, loaded with a 1:1 combination of Hsp27 and HER2 siRNAs and functionalized with RK16 and Oct, respectively; ii) FC-2s-RK16-LNPs, functionalized with the RK16 peptide and loaded with a 1:1 combination of a FAM-labeled version of Hsp27 siRNA (FAM-Hsp27 siRNA; green emission) and Cy5-HER2-siRNA; iii) C-s-Oct-LNPs, functionalized with the Oct peptide and loaded with Cy5-HER2 siRNA alone; and iv) e-RK16-LNPs and e-Oct-LNPs, functionalized with RK16 and Oct, respectively, and containing no siRNA cargo. Additional non-functionalized (naked) LNPs will serve as controls, including: i) 2s-LNPs, FC-2s-LNPs, C-s-LNPs, and scr-LNPs, loaded with the same siRNA cargoes as their respective functionalized counterparts (Hsp27 siRNA + HER2 siRNA, FAM-Hsp27 siRNA + Cy5-HER2 siRNA, Cy5-HER2 siRNA, and scrambled siRNA, respectively), and ii) e-LNPs, empty, non-functionalized LNPs. Finally, to assess the influence of PEG-lipid composition, we will prepare an additional control empty formulation, e-DSG-LNPs, in which DSG-PEG (a stearate-based PEG lipid) replaces palmitate-based DPG-PEG (Fig.1A).

**Figure 1.**
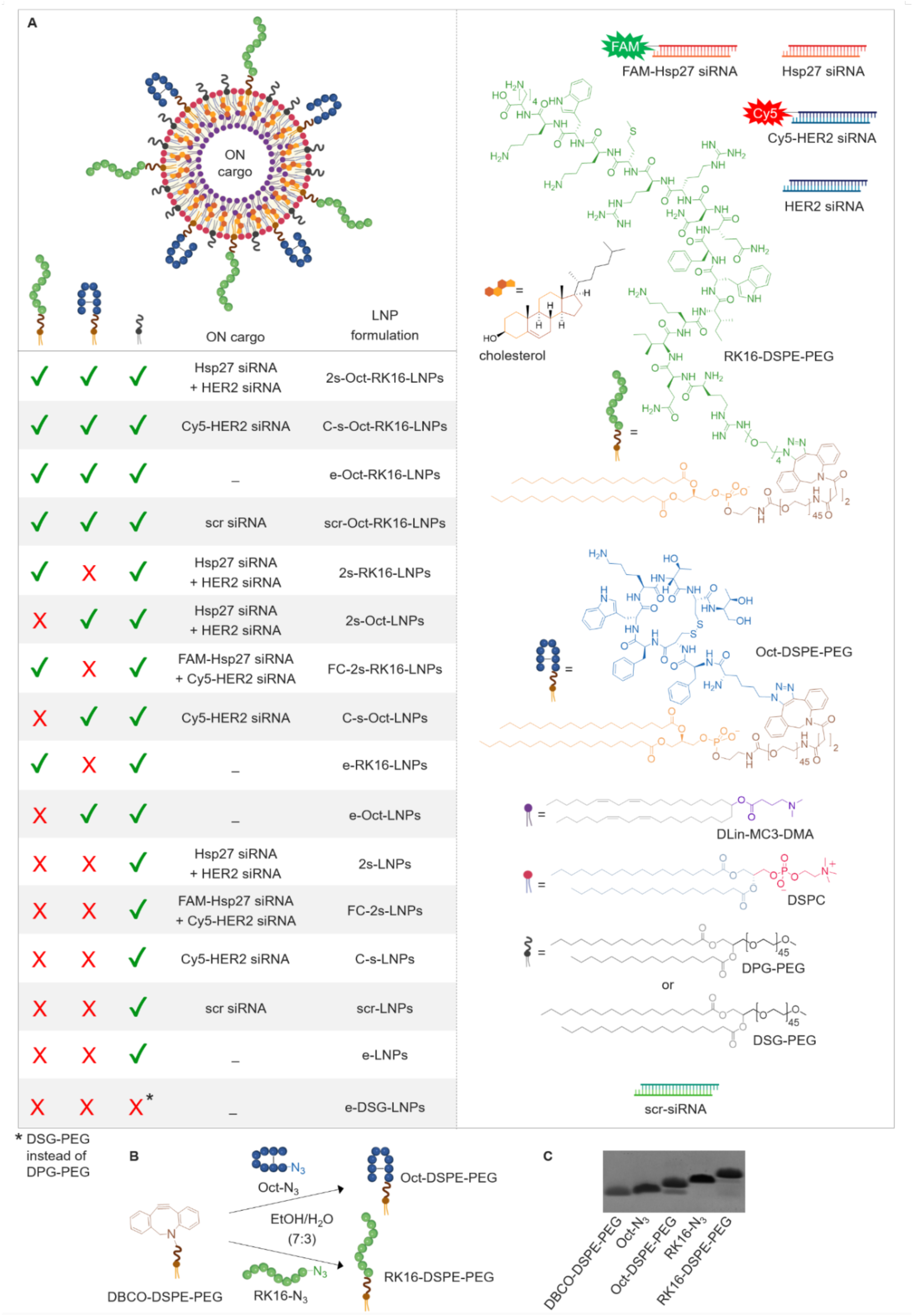
(A) Schematic representation of the LNP formulations used in this study. Formulations include dually-functionalized, singly-functionalized and non-functionalized (naked) LNPs, loaded with either: i) a combination of Hsp27 and HER2 siRNAs (FAM-and/or Cy5-labelled or unlabeled), ii) a single siRNA (Cy5-labelled HER2 siRNA or scrambled siRNA), or iii) no siRNA (empty). Structures of the peptide ligands (Oct and RK16 peptides) and the lipid components are also shown. (B) General scheme for the synthesis of Oct-DSPE-PEG and RK16-DSPE-PEG conjugates via copper-free click chemistry between azide-functionalized peptides (Oct-N_3_ and RK16-N_3_) and DBCO-functionalized DSPE-PEG. (C) Denaturing PAGE analysis of the crude conjugation reactions.

Our strategy for preparing dually modified LNPs (functionalized with both Oct and RK16) and singly-modified LNPs (functionalized with either Oct or RK16) combines copper-free click chemistry with a post-insertion approach. Ligand-conjugated DSPE-PEG lipids –Oct-DSPE-PEG and RK16-DSPE-PEG– were synthesized by reacting azido-functionalized peptides (Oct-N_3_ and RK16-N_3_) with dibenzocyclooctyne-functionalized DSPE-PEG_2000_ (DBCO-DSPE-PEG) at a 1:1 molar ratio in an EtOH/H_2_O mixture (7:3) (Fig. 1B). The efficiency of the conjugation reaction was confirmed by PAGE analysis (Fig. 1C) and gel permeation chromatography (GPC; Fig. S1), revealing conversion rates of ∼80% for Oct-DSPE-PEG and ∼95% for RK16-DSPE-PEG. Successful formation of the peptide-lipid conjugates was further validated by a reduction in absorbance at the DBCO-specific excitation wavelength (309 nm; Fig. S2) and by MALDI-TOF mass spectrometry of the crude products (Fig. S3).

The resulting Oct-DSPE-PEG and RK16-DSPE-PEG conjugates were then post-inserted into preformed LNPs –2s-LNPs, C-s-LNPs, e-LNPs, or scr-LNPs (see Methods for details)– yielding dually functionalized formulations: 2s-Oct-RK16-LNPs, C-s-Oct-RK16-LNPs, e-Oct-RK16-LNPs, and scr-Oct-RK16-LNPs (Fig. 1A). Using the same post-insertion method but using a single peptide-lipid conjugate (Oct-DSPE-PEG or RK16-DSPE-PEG), we also generated the singly-functionalized formulations 2s-Oct-LNPs, 2s-RK16-LNPs, C-s-Oct-LNPs, FC-2s-RK16-LNPs, e-Oct-LNPs and e-RK16-LNPs (Fig. 1A).

### 2.2. LNP characterization

The physicochemical properties of LNPs loaded with a 1:1 mixture of Hsp27 and HER2 siRNAs are summarized in Table 1. Functionalized LNPs exhibited increased average hydrodynamic diameters compared to non-functionalized (naked) LNPs. Specifically, the sizes of 2s-RK16-LNPs, 2s-Oct-LNPs and 2s-Oct-RK16-LNPs were 85.61 ± 0.72 nm, 110.40 ± 0.87 nm, and 119.00 ± 0.96 nm, compared to 70.36 ± 1.22 nm for the naked 2s-LNPs. Correspondingly, the polydispersity index (PDI) increased in the functionalized systems (0.250, 0.290, and 0.250 for 2s-RK16-LNPs, 2s-Oct-LNPs, and 2s-Oct-RK16-LNPs, respectively) versus a PDI of 0.190 for 2s-LNPs.

**Table 1.** Physicochemical characterization of non-decorated and decorated LNPs.

| LNPs | DLS<br>hydrodynamic<br>diameter (nm) | Polydispersity<br>index | Zeta potential<br>(mV) | Encapsulation<br>efficiency |
| --- | --- | --- | --- | --- |
| 2s-LNPs | 70.36 ± 1.22 | 0.190 | -2.77 ± 0.99 | 83.9 % |
| 2s-RK16-LNPs | 85.61 ± 0.72 | 0.250 | +9.29 ± 0.38 | 93.8 % |
| 2s-Oct-LNPs | 110.40 ± 0.87 | 0.290 | -3.15 ± 1.87 | 82.1 % |
| 2s-RK16-Oct-LNPs | 119.00 ± 0.96 | 0.250 | -1.94 ± 0.57 | 79.8 % |

Zeta potential (ZP) measurements reflected the influence of surface ligand charge. The 2s-RK16-LNPs displayed a significantly higher positive ZP (+9.29 ± 0.38 mV) relative to the 2s-LNPs, which exhibited a slightly negative ZP (−2.77 ± 0.99 mV). The ZP of 2s-Oct-LNPs was similar to that of the naked system (−3.15 ± 1.87 mV). This increase in surface charge with RK16 functionalization is consistent with the strong positive net charge of RK16-DSPE-PEG lipid (+6), compared to the neutral or mildly positive charges associated with DPG-PEG (0) and Oct-DSPE-PEG (+1), respectively. As expected, the ZP of the dually-functionalized 2s-Oct-RK16-LNPs (−1.94 ± 0.57 mV) fell between those of the of the singly-functionalized counterparts.

Encapsulation efficiency (EE) was slightly reduced in the functionalized systems. The EE of 2s-Oct-RK16-LNPs was 79.8 %, marginally lower than the 83.9 % observed for naked 2s-LNPs, likely due to siRNA loss during ligand insertion. The apparently higher encapsulation efficiency observed with RK16-decorated LNPs (93.8%) was considered an unavoidable artifact, likely caused by electrostatic interactions between the abundant cationic CPP and free nucleic acids, which may interfere with fluorimetric quantitation.

Both naked and dually-functionalized LNPs loaded with a 1:1 mixture of Hs27 siRNA and HER2 siRNAs (2s-LNPs and 2s-Oct-RK16-LNPs, respectively) demonstrated excellent colloidal stability over a seven-day period when stored at 4 °C (Fig. S4). Repeated measurements confirmed that key physicochemical properties –including average size, hydrodynamic diameter, surface charge, and encapsulation efficiency–remained stable throughout the storage period for both incubations. Notably, the initial differences observed between the two LNP types (e.g., size and surface charge) were maintained consistently over time (Fig. S4 A-D).

### 2.3. Synergistic antiproliferative effects of combined Hsp27 and HER2 siRNA treatment

Building on previous evidence that dual inhibition of Hsp27 and HER2 –achieved using a small molecule inhibitor and an anti-HER2 antibody– elicits a synergistic antiproliferative effect in HER2+ BC cells,^46^ we next assessed the efficacy of co-delivering Hsp27 and HER2 siRNAs using our naked LNP formulation (2s-LNP formulation). This formulation, encapsulating a 1:1 molar ratio of the two siRNAs, was tested in HER2+ SK-BR-3 (Fig. 2A) and BT-4T4 (Fig. 2B) cell lines. In both models, the combination treatment significantly inhibited cell proliferation in a synergistic manner, as evidenced by isobologram analyses with combination index (CI) values of 0.74 for SK-BR-3 (Fig. 2A) and 0.68 for BT-474 (Fig. 2B) cells after 96 hours of treatment.

**Figure 2.**
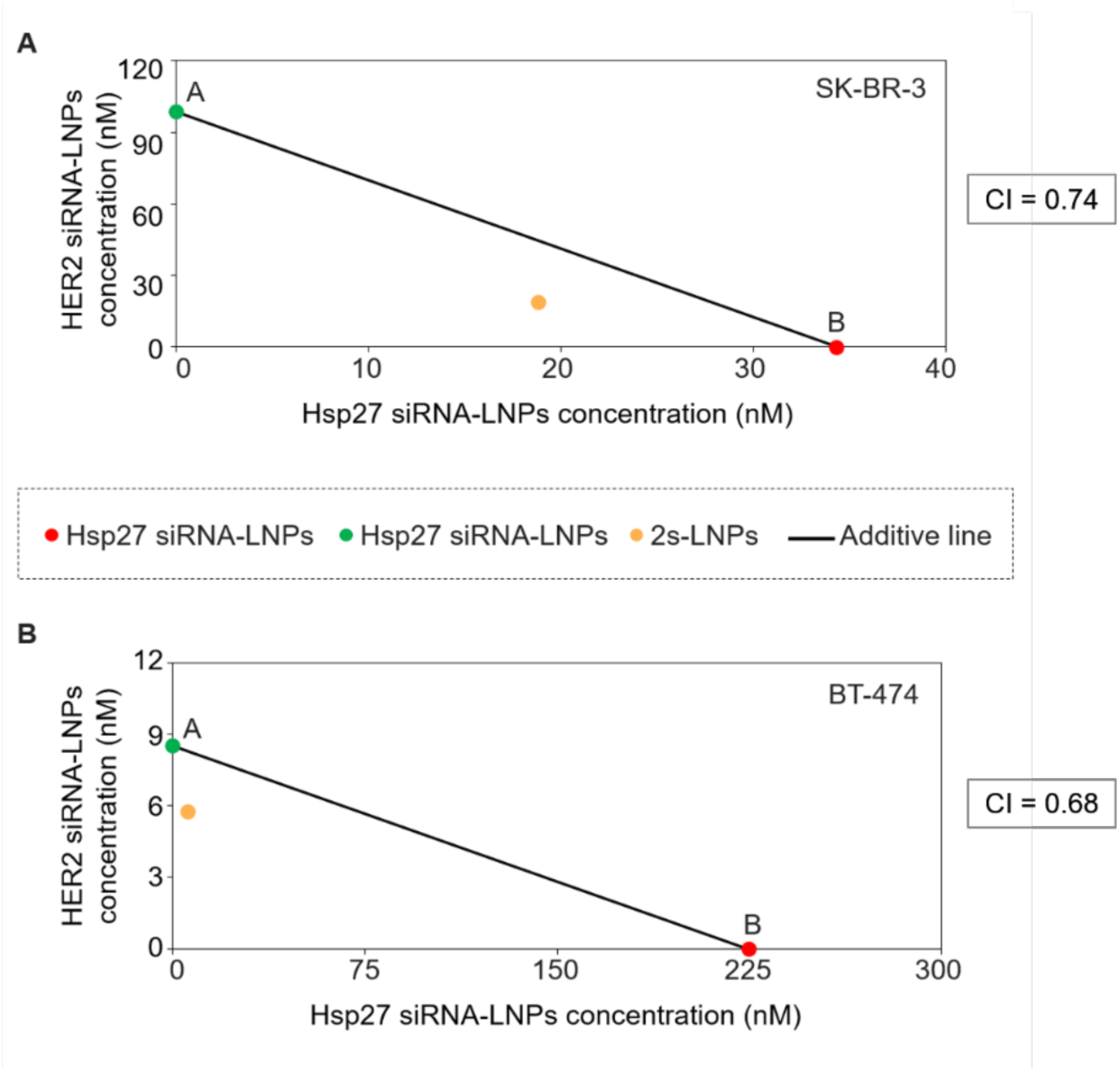
Isobologram of naked LNPs loaded with individual siRNAs (either Hsp27 siRNA or HER2 siRNA) in SK-BR-3 (A) and BT-474 (B) cells. The IC_50_ additive isobole is a straight line between points A and B. A and B indicate the IC_50_ values of HER2 siRNA-loaded LNPs and Hsp27 siRNA-loaded LNPs, respectively. The yellow-colored dot represents the IC_50_ value of the 2s-LNPs treatment (naked LNPs loaded with a 1:1 mixture of Hsp7 and HER2 siRNAs). The area above the straight line indicates antagonism and the area below the additive isobole suggests synergism. CI: combination index.

### 2.4. Evaluating individual contributions of LNP decorated molecules

#### 2.4.1. RK16 cell penetrating functionality

We next investigated the cell penetrating capacity of the RK16 ligand in SK-BR-3 and BT-474 cell lines by confocal microscopy. To this end, we used singly-modified LNPs functionalized with RK16 and co-loaded with FAM-Hsp27 and Cy5-HER2 siRNAs (FC-2s-RK16-LNPs). As controls, naked LNPs loaded with the same fluorescent siRNA combination (FC-2s-LNPs) were used.

In SK-BR-3 cells, the internalization of FC-2s-RK16-LNPs was markedly higher than that observed for naked FC-2s-LNPs, as evidenced by stronger red (Cy5) and green (FAM) fluorescent signals (Fig. 3A) and confirmed by quantitative analysis (Fig. 3C). At a 20 nM siRNA dose, FC-2s-RK16-LNPs efficiently entered cells within 24 hours, while naked FC-2s-LNPs showed negligible uptake. Although delayed red and green fluorescence signals were detected in FC-2s-LNP-treated cells after an additional 24 hours (Fig. S5A and C), the signal intensity remained approximately fivefold lower than that observed with FC-2s-RK16-LNPs.

**Figure 3.**
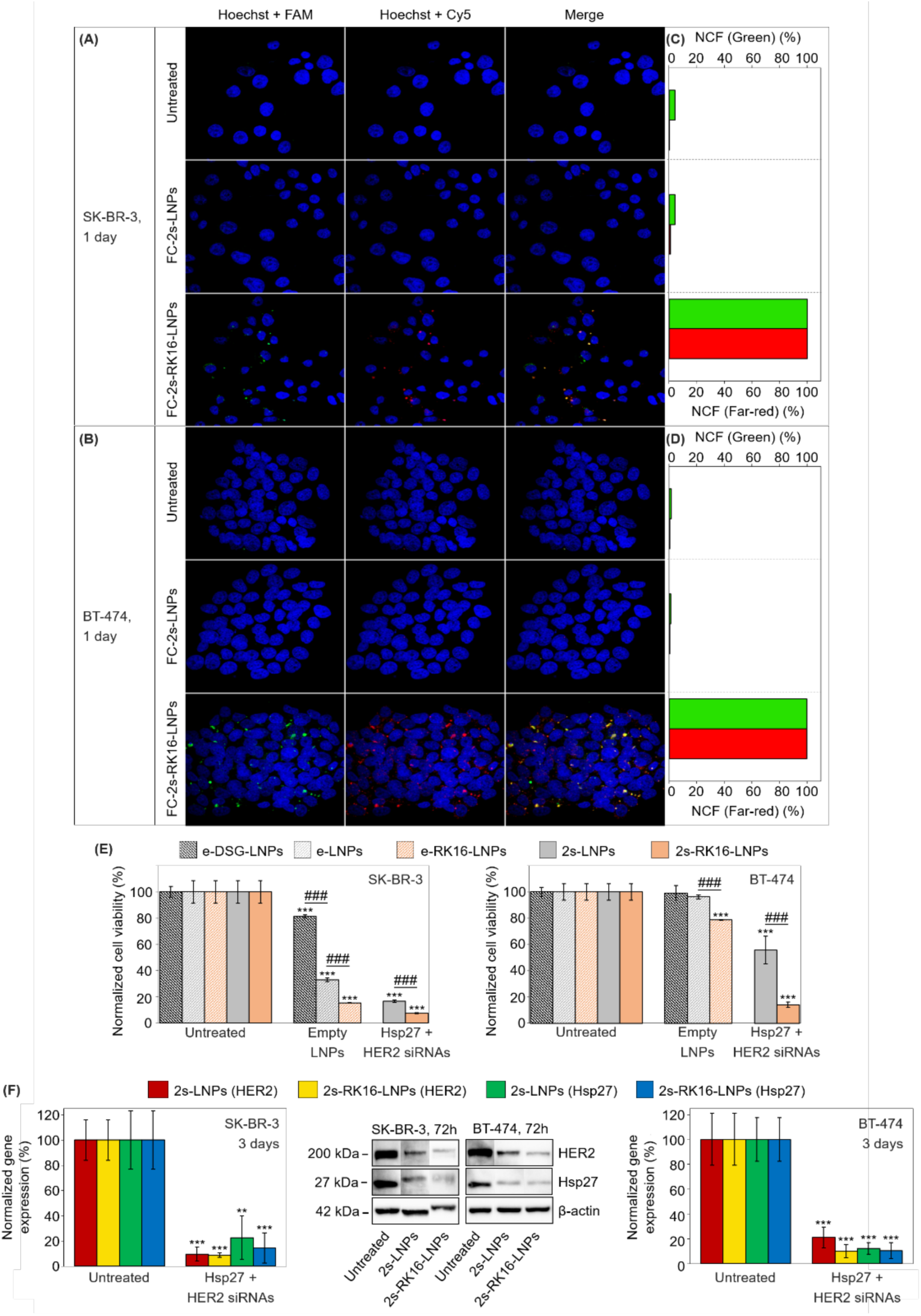
(A, B) Confocal microscopy images of SK-BR-3 (A) and BT-474 (B) cells incubated for 24 hours with either naked or RK16-functionalized LNPs, both loaded with a 1:1 mixture of FAM-Hsp27 and Cy5-HER2 siRNAs (FC-2sLNPs and FC-2s-RK16-LNPs, respectively). Untreated cells served as negative controls. (C, D) Quantification of green (FAM) and red (Cy5) fluorescence intensities from images shown in panels (A) and (B). (E) Crystal violet cell viability assay performed 96 hours post-transfection of SK-BR-3 and BT-474 cells with either naked or RK16-functionalized LNPs, which were either empty or loaded with a 1:1 mixture of Hsp27 and HER2 siRNAs (e-LNPs, e-RK16-LNPs, 2s-LNPs, and 2s-RK16-LNPs). Naked empty LNPs formulated with DSG-PEG instead of DPG-PEG (e-DSG-LNPs) were also used as controls. (F) Representative immunoblots and quantitative analysis of protein expression for HER2, Hsp27 and actin (internal control) from the same cell lines described in panel (E) treated with 2s-LNPs and 2s-RK16-LNPs for 72 hours. In all experiments (panels A-F), the total siRNA concentration was 20 nM. Results in panels (E) and (F) were normalized to untreated controls. Independent experiments were performed and quantified (n = 3 for both Western blot and viability assays). Data are expressed as mean ± standard deviation. For panels (E) and (F), unpaired Student’s *t*-tests were used to compare each sample to the untreated control and between selected samples. Symbols: ** p < 0.01; ### / *** p < 0.001. Asterisks (*) denote significance versus untreated control; hash symbols (#) indicate significance versus experimental groups.

Similar trends were observed in BT-474 cells. RK16-functionalized FC-2s-RK16-LNPs demonstrated superior intracellular delivery at both time points (24 and 48 hours) and at the same siRNA concentration (20 nM), compared to naked FC-2s-LNPs (Fig. 3 B, D and Fig. S5B, D). These results confirm that RK16 enhances cellular uptake through its cell penetrating peptide activity, promoting efficient delivery of siRNA cargo into HER2+ BC cells.

Having confirmed the RK16-mediated enhancement of intracellular delivery, we next examined whether this translated into improved biological activity. Before assessing antiproliferative effects of the siRNA-loaded RK16-LNPs, we evaluated whether the lipid composition of the LNPs themselves influenced cell viability. Previous studies have shown that HER2+ BC cells depend heavily on fatty acid synthesis for survival,^53^ and that disrupting this pathway can inhibit growth and induce apoptosis. In particular, palmitate supplementation has been linked to key metabolic changes –such as AMPK activation and fatty acid synthesis inhibition– that impact glycolysis and glutamine metabolism. This lipotoxicity appears to be more pronounced in certain HER2+ BC cell lines, like SK-BR-3, compared to others like BT-474, possibly due to intrinsic genetic differences and compensatory mechanisms in the latter.

Given the presence of palmitate in our LNPs (via DPG-PEG), we compared the effects of empty naked LNPs formulated with either palmitate-based DPG-PEG (e-LNPs) or stearate-based DSG-PEG (e-DSG-LNPs) on cell proliferation. In SK-BR-3 cells, palmitate-based e-LNPs reduced cell proliferation by (67 ± 1) % after 96 hours, whereas e-DSG-LNPs led to only a (19 ± 1) % reduction (Fig. 3E). In contrast, BT-474 cells were largely unaffected by palmitate-based LNPs, exhibiting only a minimal growth inhibition [(4 ± 1) %] in response to e-LNPs, which was comparable to the effect caused by e-DSG-LNPs in this cell line [(1 ± 5) % growth inhibition] (Fig. 3E). These results are consistent with previous findings^53^ and suggest that the palmitate moiety in DPG-PEG contributes selectively to cytotoxicity in palmitate-sensitive cell lines like SK-BR-3 cells, which may offer an added therapeutic advantage when targeting specific tumor subtypes. These findings underscore the need for further investigation into the molecular mechanisms underlying cancer cell sensitivity to certain metabolites. Together with the recent development of genetic tests enabling the analysis of gene expression clusters and a more refined molecular subtyping of BC, the adaptable composition of LNPs offers a promising strategy for the personalized treatment of HER2+ BC and potentially other malignancies by tailoring formulations to individual tumor profiles.

The RK16 ligand itself induced additional toxicity beyond that observed for the naked LNPs [(85 ± 0.1) % and (21 ± 0.1) % growth inhibition for e-RK16-LNP-treated SK-BR-3 and BT-474 cells versus (67 ± 1) % and (4 ± 1) % inhibition for the same cell lines treated with e-LNPs, respectively; Fig. 3E]. In any case, inclusion of the 2siRNA cargo (Hsp27 + HER2 siRNAs) in 2s-RK16-LNPs further amplified the cytotoxic effect of the LNPs. Thus, treatment of SK-BR-3 and BT-4T4 cells with 2s-RK16-LNPs led to a marked reduction in cell proliferation relative to both empty LNPs [naked (e-LNPs) or RK16-functionalized (e-RK16-LNPs)] and naked siRNA-loaded 2s-LNPs. In SK-BR-3 cells treated with 2s-RK16-LNPs (20 nM total siRNA), proliferation was reduced by (93 ± 0.3) %, compared to (83 ± 1) % with 2s-LNPs. A similar trend was observed in BT-474 cells, with growth inhibition of (86 ± 2) % for 2s-RK16-LNPs and (44 ± 10) % for 2s-LNPs.

Western blot (WB) analysis confirmed that the increase in cytotoxicity observed with 2siRNA-loaded LNPs arose from the simultaneous silencing of Hsp27 and HER2 genes (Fig. 3F). Treatment with 2s-RK16-LNPs led to strong gene knockdown, with (86 ± 16) % and (91 ± 2) % reductions in Hsp27 and HER2 expression, respectively, in SK-BR-3 cells, and (89 ± 6) % and (90 ± 5) % knockdown in BT-474 cells. Although these knockdown levels were comparable to those achieved with naked 2s-LNPs [(77 ± 17) % and (90 ± 6) % in SK-BR-3 cells; (88 ± 5) % and (79 ± 8) % in BT-474 cells], the 2siRNA-loaded RK16-functionalized LNPs produced superior cytotoxic effects (Fig. 3E). This enhanced efficacy is likely due to more efficient uptake across a broader population of cells within a shorter time frame.

#### 2.4.2. Octreotide functionality

After confirming the cell penetrating capability of the RK16 ligand, we next assessed the cell-selective targeting properties conferred by the Oct functionality. Specifically, we evaluated whether singly-modified LNPs functionalized with Oct could selectively recognize and preferentially internalize into tumor cells overexpressing the SSTR2 receptor in the presence of other (non-overexpressing) cell types. Co-culture models are well-established platforms for assessing the targeting selectivity of ligand-decorated delivery systems.^54^ For this purpose, a parental cell line with low-to moderate receptor expression is co-cultured with a fluorescently labeled derivative engineered to overexpress the same receptor, allowing for direct assessment of ligand-mediated uptake –using a distinguishable fluorophore– via microscopy or flow cytometry.

To establish such models, we generated SK-BR-3 SSTR2OE (GFP+) and BT-474 SSTR2OE (GFP+) cell lines through viral transduction (see Methods and Fig. S6), achieving stable overexpression of both SSTR2 and green fluorescent protein (GFP). Flow cytometry analysis confirmed elevated SSTR2 surface expression in the modified cells (Fig. S6A-E), with mean fluorescence intensities of 55.20 and 16.4, respectively, compared to 2.28 and 2.17 in their parental lines (SK-BR-3 and BT-474, respectively). GFP expression was also verified via both flow cytometry and fluorescence microscopy (Fig. S6A-F).

To assess octreotide-mediated targeting, we treated co-cultures of parental and SSTR2-overexpressing (SSTR2OE) (GFP+) cells [SK-BR-3 + SK-BR-3 SSTROE (GFP+) and BT-474 + BT-474 SSTROE (GFP+) co-cultures] with Cy5-labelled HER2 siRNA-loaded Oct-functionalized LNPs (C-s-Oct-LNPs) for 24 h. Control groups of cells were treated with non-targeted LNPs loaded with the same Cy5-labeled siRNA (C-s-LNPs). Confocal microscopy (Fig. 4A, B) and quantitative fluorescence analysis (Fig. 4C, D) revealed pronounced accumulation of red fluorescence (Cy5-HER2 siRNA) in GFP+ SSTR2-overexpressing cells, both in SK-BR-3 (Fig. 4A, C) and BT-474 (Fig. 4B, D) co-cultures treated with C-s-Oct-LNPs. Cy5 signal was predominantly localized within the cytoplasm of GFP+ cells, demonstrating selective uptake of C-s-Oct-LNPs. In contrast, no such cell selectivity was observed with non-targeted C-s-LNPs, where red fluorescence was distributed randomly across both GFP+ and GFP-populations.

**Figure 4.**
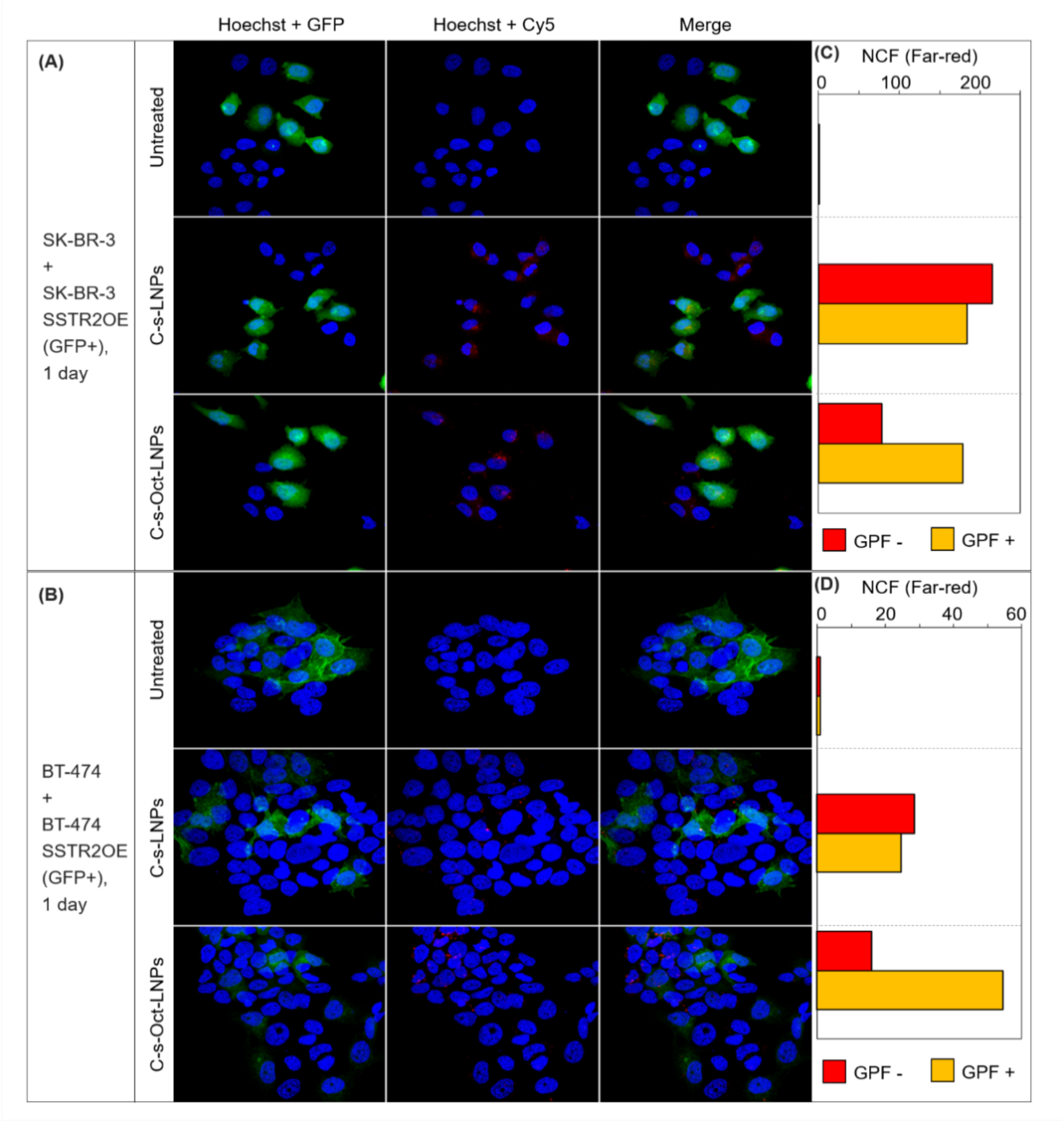
(A, B) Confocal microscopy images of co-cultures consisting of SK-BR-3 and SK-BR-3 SSTR2OE (GFP+) cells (A), and BT-474 and BT-474 SSTR2OE (GFP+) cells (B), incubated for 24 hours with either naked or Oct-functionalized LNPs, both loaded with Cy5-labeled HER2 siRNA (C-s-LNPs and C-s-Oct-LNPs, respectively) at a final siRNA concentration of 40 nM. Untreated cells served as negative controls. (C, D) Quantification of red (Cy5) fluorescence intensity in GFP-cells (red) and GFP+ cells (yellow, indicating co-localization of Cy5 and GFP signals) from images shown in panels (A) and (B).

However, the selective accumulation of C-s-Oct-LNPs in SSTR2OE (GFP+) cells was less pronounced at 48 hours post-transfection in SK-BR-3 + SK-BR-3 SSTR2OE (GFP+) and BT-474 + BT-474 SSTR2OE (GFP+) co-cultures, with only modest enrichment of Cy5-labeled siRNA observed in the SSTR2OE GFP+ population (Fig. S7). This apparent decline in accumulation is likely showing the early death of the SSTR2OE cells, potentially triggered by increased internalization of HER2-targeting siRNA delivered by the Oct-functionalized LNPs. Overall, the data clearly demonstrate that Oct-decorated LNPs achieve preferential siRNA delivery to SSTR2-overexpressing tumor cells during the early phase post-treatment in mixed populations.

Octreotide alone has been reported to exert antiproliferative effects in both endocrine and BC cells, though these effects vary depending on dose, exposure time, and cell type.^55^ Based on these observations and the selective uptake demonstrated in our previous internalization studies (Fig. 4), we next evaluated the potential cytotoxicity of Oct-functionalized LNPs in monocultures and co-cultures of HER2+ BC cells. Crystal violet viability assays were performed on SK-BR-3, SK-BR-3 SSTR2OE (GFP+), BT-474, and BT-474 SSTR2OE (GFP+) monocultures treated with empty naked LNPs (e-LNPs) or Oct-decorated LNPs (e-Oct-LNPs) (Fig. S8). Cell viability was measured 96 hours post-treatment to assess the influence of Oct-functionalization and SSTR2 expression. Overall, both LNP formulations induced similar levels of cytotoxicity across all cell types. Only a modest increase in cell death was observed in BT-474 SSTR2OE cells treated with e-Oct-LNPs [(15 ± 4) % inhibition of cell proliferation], compared to the same cell line treated with naked empty e-LNPs [(3 ± 5) % inhibition], suggesting limited Oct-mediated cytotoxicity under these conditions. Consistent with our earlier observations, SK-BR-3 and SK-BR-3 SSTR2OE cells were highly sensitive to lipid exposure, displaying reduced viability, regardless of Oct functionalization [(70 ± 1) % and (65 ± 1) % growth inhibition for SK-BR-3 cells treated with e-LNPs and e-Oct-LNPs, and (64 ± 1) % and (58 ± 1) % inhibition for SK-BR-3 SSTROE (GFP+) cells analogously treated].

To validate the selective targeting observed in uptake assays (Fig. 4), we next conducted flow cytometry viability studies in 1:1 co-cultures of SK-BR-3 + SK-BR-3 SSTR2OE (GFP+) and BT-474 + BT-474 SSTR2OE (GFP+) cells (Fig. 5). These co-cultures were treated with naked or Oct-functionalized LNPs loaded with a 1:1 mixture of Hsp27 and HER2 siRNAs (2s-LNPs and 2s-Oct-LNPs, respectively). Viability studies performed at 48 hours (in the case of SK-BR-3 + SK-BR-3 SSTR2OE co-cultures) and 72 hours (in the case of BT-474 + BT-474 SSTR2OE co-cultures) post-treatment revealed comparable overall cytotoxicity between formulations (Fig. 5A), with global cell death rates of (50 ± 20) % and (45 ± 10) % in SK-BR-3 + SK-BR-3 SSTR2OE co-cultures after treatment with 2s-LNPs and2s-Oct-LNPs, respectively. Similarly, global cell death rates of (42 ± 7) % and (37 ± 4) % were observed in BT-474 + BT-474 SSTR2OE co-cultures. Remarkably, Oct-functionalized formulations (2s-Oct-LNPs) selectively reduced the proportion of GFP+ (SSTR2-overexpressing) cells within both co-culture population. Specifically, the percentage of GFP+ cells decreased from (49 ± 2) % and (49 ± 4) % in the untreated SK-BR-3 and BT-474 co-cultures, respectively, to (43 ± 10) % and (39 ± 4) % following treatment (Fig. 5B). This effect that was not observed with non-targeted (naked) LNPs. The reduction in GFP+ cell representation strongly supports the receptor-mediated selectivity of the Oct-functionalized delivery system, confirming its preferential targeting within heterogeneous tumor cell populations.

**Figure 5.**
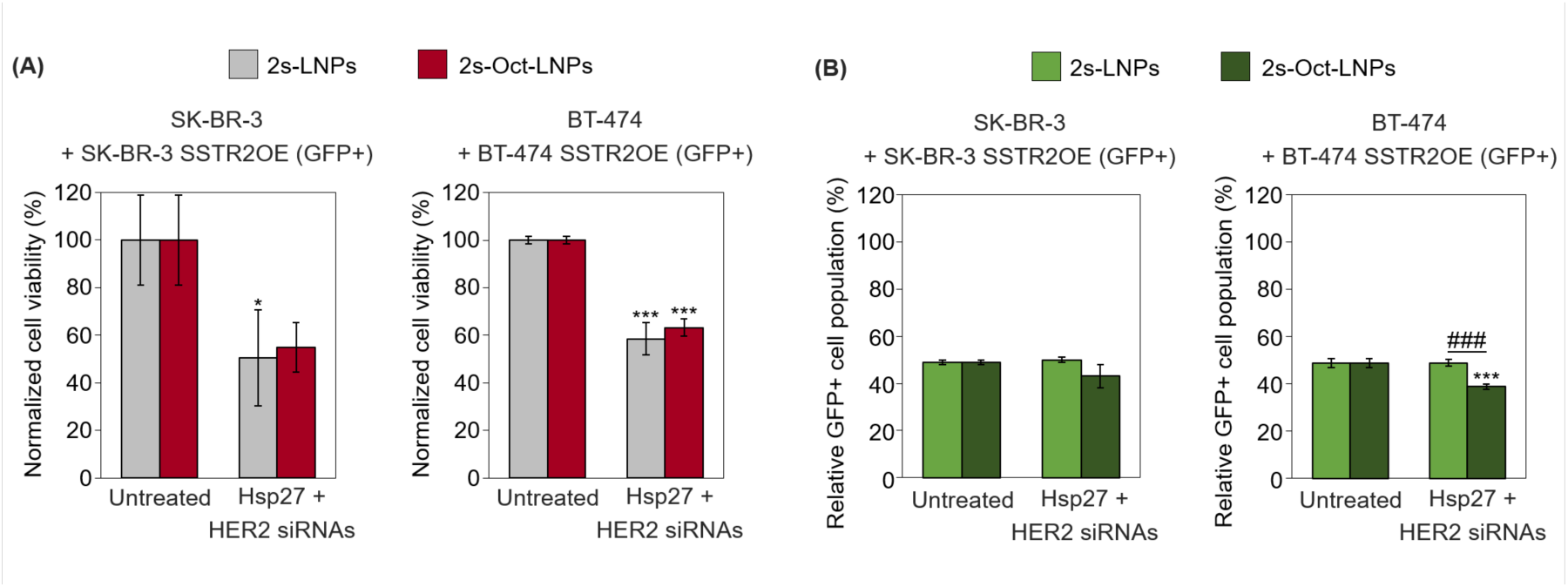
(A) Flow cytometric cell viability assay of SK-BR-3 + SK-BR-3 SSTR2OE (GFP+) and BT-474 + BT-474 SSTR2OE (GFP+) co-cultures treated with either naked or Oct-functionalized LNPs, both loaded with a 1:1 mixture of Hsp27 and HER2 siRNAs (2s-LNPs and 2s-Oct-LNPs) at a final siRNA concentration of 20 nM. Assays were performed 48 hours post-transfection for SK-BR-3 co-cultures and 72 hours for BT-474 co-cultures. (B) Flow cytometric quantification of the relative abundance of viable GFP+ cells from the co-cultures described in panel (A). Results were normalized to untreated controls. Independent experiments were performed and quantified (n = 3). Results are represented as mean ± standard deviation. Statistical comparisons were performed using unpaired Student’s *t*-tests. Symbols: * p < 0.05; *** / ### p < 0.001*. Asterisks (*) denote significance versus untreated control; hash symbols (#) indicate significance versus experimental groups.

### 2.5. Dually-functionalized LNPs

After confirming the enhanced delivery efficiency of the cell-penetrating peptide RK16 and the SSTR2 selectivity conferred by the receptor-targeting peptide octreotide, we evaluated their combined effects in dually-functionalized LNPs (Oct-RK16-LNPs).

To determine whether the improved performance previously observed with RK16-decorated LNPs was retained upon dual functionalization, we performed crystal violet viability assays on SK-BR-3, BT-474, SK-BR-3 SSTR2OE (GFP+), and BT-474 SSTR2OE (GFP+) monocultures 96 hours post-transfection (Fig. 6A). Cells treated with empty dually-decorated LNPs (e-Oct-RK16-LNPs) displayed cytotoxic profiles similar to those of empty naked (e-LNPs) and RK16-decorated (e-RK16-LNPs) formulations (see Fig. 3E for e-RK16-LNPs). Toxicity remained mild in BT-474 and BT-474 SSTR2OE (GFP+) cells (death rates < 15%), while SK-BR-3 and SK-BR-3 SSTR2OE cells showed pronounced sensitivity, with (87 ± 1) % and (83 ± 2) % cell death, respectively –lower than with siRNA-loaded counterparts (2s-Oct-RK16-LNPs), but still notable. Remarkably, dually decorated LNPs co-loaded with a 1:1 mixture of Hsp27 and HER2 siRNAs (2s-Oct-RK16-LNPs) outperformed their naked 2siRNA-loaded counterparts (2s-LNPs), inducing significantly greater reductions in cell viability across all tested cell lines. At a total siRNA dose of 20 nM, 2s-Oct-RK16-LNPs induced (94 ± 0.3) %, (92 ± 0.4) %, (67 ± 6) %, and (69 ± 1) % cell death rates in SK-BR-3, SK-BR-3 SSTROE (GFP+), BT-474 and BT-474 SSTR2OE (GFP+) cells, respectively, compared to (87 ± 1) %, (85 ± 2) %, (37 ± 8) % and (43 ± 5) % with naked 2s-LNP (Fig. 6A).

**Figure 6.**
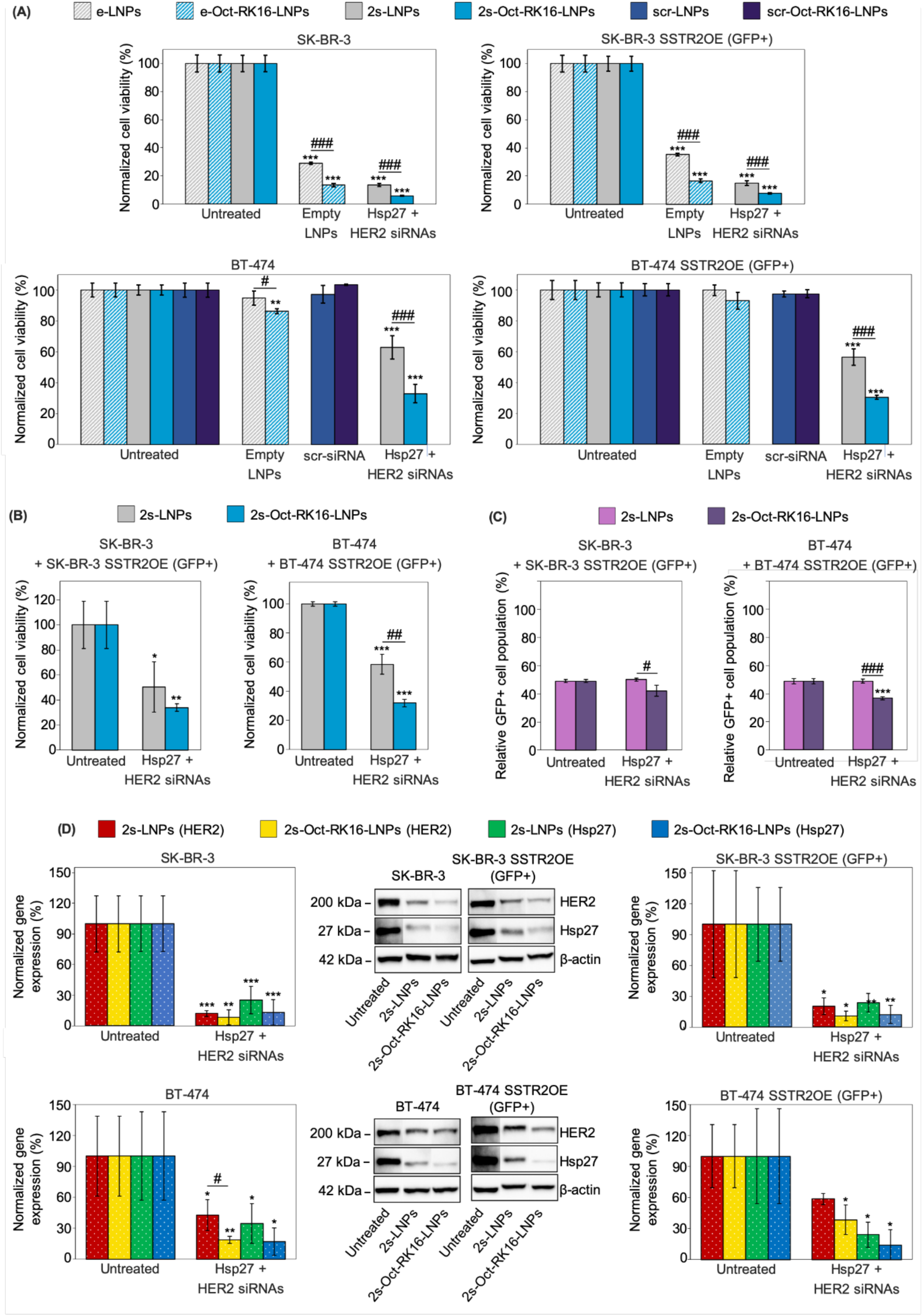
(A) Crystal violet cell viability assay performed 96 hours post-transfection of SK-BR-3, SK-BR-3 SSTROE (GFP+), BT-474, and BT-474 SSTR2OE (GFP+) cells. Cells were treated with either naked or dually-functionalized (Oct-RK16) LNPs, which were either empty or loaded with a 1:1 mixture of Hsp27 and HER2 siRNAs, or with scrambled siRNA (e-LNPs, e-Oct-RK16-LNPs, 2s-LNPs, 2s-Oct-RK16-LNPs, scr-LNPs, and scr-Oct-RK16-LNPs). (B) Flow cytometric cell viability assay of SK-BR-3 + SK-BR-3 SSTR2OE (GFP+) and BT-474 + BT-474 SSTR2OE (GFP+) co-cultures treated with either naked or dually-functionalized LNPs, both loaded with a 1:1 mixture of Hsp27 and HER2 siRNAs (2s-LNPs and 2s-Oct-RK16-LNPs). Assays were performed 48 hours post-transfection for SK-BR-3 co-cultures and 72 hours for BT-474 co-cultures. (C) Flow cytometric quantification of the relative abundance of viable GFP+ cells in the co-cultures described in panel (B). (D) Representative immunoblots and quantitative analysis of protein expression for HER2, Hsp27 and actin (internal control) from the same cell lines described in panel (A) treated with 2s-LNPs, and 2s-Oct-RK16-LNPs for 72 hours. In all experiments (panels (A-D), the total siRNA concentration was 20 nM. Results were normalized to untreated controls. Independent experiments were performed and quantified (n = 3 for both viability and Western blot assays). Data are expressed as mean ± standard deviation. Unpaired Student’s *t*-tests were used to compare each sample to the untreated control and between selected samples. Symbols: * / # p < 0.05; ** / ## p < 0.01; *** / ### p < 0.001. Asterisks (*) denote significance versus untreated control; hash symbols (#) indicate significance versus experimental groups.

Additionally, both naked and dually-functionalized LNPs loaded with scrambled siRNA (scr-LNPs and scr-Oct-RK16-LNPs) performed similarly to their empty counterparts (e-LNPs and e-Oct-RK16-LNPs), confirming that the observed cytotoxicity was siRNA-specific. Importantly, no significant cytotoxicity was observed in the non-tumorigenic breast epithelial cell line MCF-10A following treatment with either empty (e-LNPs, e-Oct-RK16-LNPs) or 2siRNA-loaded (2s-LNPs and 2s-Oct-RK16-LNPs) formulations (Fig. S9). Cell death remained below 20 % at a 20 nM siRNA dose, supporting the safety and tolerability of these formulations in normal cells and their potential for *in vivo* application.

In summary, 2siRNA-loaded, dually-decorated LNPs significantly reduced cancer cell viability and demonstrated enhanced potency over naked LNPs, comparable to RK16-decorated LNPs. The reduced surface charge of these formulations may further improve their therapeutic potential by enhancing pharmacokinetic profiles and minimizing immunogenicity *in vivo*.

We next evaluated whether the SSTR2 selectivity previously observed with Oct-decorated LNPs was retained in the dually functionalized Oct-RK16 systems. To this end, we conducted flow cytometry viability assays using 1:1 co-cultures of SK-BR-3 with SK-BR-3 SSTR2OE (GFP+) cells and BT-474 with BT-474 SSTR2OE (GFP+) cells, each transfected with either 2s-LNPs or 2s-Oct-RK16-LNPs (Fig. 6B, C). Analysis of GFP+ cell abundance in these co-cultures revealed that dual peptide decoration did not compromise SSTR2 selectivity, despite the unspecific potency enhancement attributed to RK16. While total cell death was (50 ± 20) % and (41 ± 7) % in SK-BR-3 and BT-474 co-cultures treated with 2s-LNPs, it rose to (66 ± 3) % and (68 ± 3) %, respectively, after treatment with 2s-Oct-RK16-LNPs (Fig. 6B). Moreover, the proportion of GFP+ cells decreased from (49 ± 2) % and (49 ± 4) % in the untreated SK-BR-3 and BT-474 co-cultures, to (42 ± 3) % and (37 ± 3) % following treatment with the dually-decorated 2s-Oct-RK16-LNPs (Fig. 6C), a decrease that was not observed with the naked formulation (2s-LNPs). These results indicate that RK16 and Oct retain their individual functionalities when combined, contributing complementary effects to the overall improved cytotoxic response.

Western blot analysis (Fig. 6D) confirmed robust inhibition of Hsp27 and HER2 expression in SK-BR-3, SK-BR-3 SSTR2OE, BT-474 and BT-474 SSTR2OE monocultures following treatment with 2s-Oct-RK16-LNPs, comparable to –and in the case of HER2 expression in BT-474 cells, slightly greater than– those achieved with naked 2s-LNPs. In all cases, protein expression was consistently reduced by at least 70 % at a total siRNA dose of 20 nM. In contrast, scrambled siRNA-loaded naked (scr-LNPs) and dually decorated LNPs (scr-Oct-RK16-LNPs) did not significantly affect target protein levels (Fig. S10), confirming the specificity of the observed effects. Overall, our results confirm that the dually-decorated 2siRNA-loaded LNP formulation induces superior anti-proliferative effects and specificity compared to non-functionalized versions, with RK16 and Oct maintaining their cell-penetrating and targeting abilities.

Finally, to assess the performance of naked and dually-decorated LNPs under conditions that more closely mimic physiological environments, cell uptake studies were conducted using 1:1 co-cultures of MCF-10A cells with either SK-BR-3 SSTR2OE (GFP+) or BT-474 SSTR2OE (GFP+) cells. These co-cultures simulate the heterogeneous tumor microenvironment commonly observed in patients, such as HER2+ BC tumors surrounded by non-tumorigenic breast epithelial cells. Both co-culture systems were transfected with Cy5-HER2 siRNA-loaded naked or dually-functionalized LNPs (C-s-LNPs, C-s-Oct-RK16-LNPs).

Confocal microscopy analysis performed 24 hours post-transfection revealed that dually-functionalized LNPs demonstrated higher internalization efficiency compared to naked LNPs, as evidenced by increased Cy5 fluorescence intensity (Fig. 7A, C and Fig. S10A, C). Furthermore, C-s-Oct-RK16-LNPs preferentially accumulated in SSTR2-overexpressing (GFP+) cells [SK-BR-3 (Fig. 7A, C) or BT-474 (Fig.S10A, C)], with minimal uptake observed in surrounding non-tumor cells.

**Figure 7.**
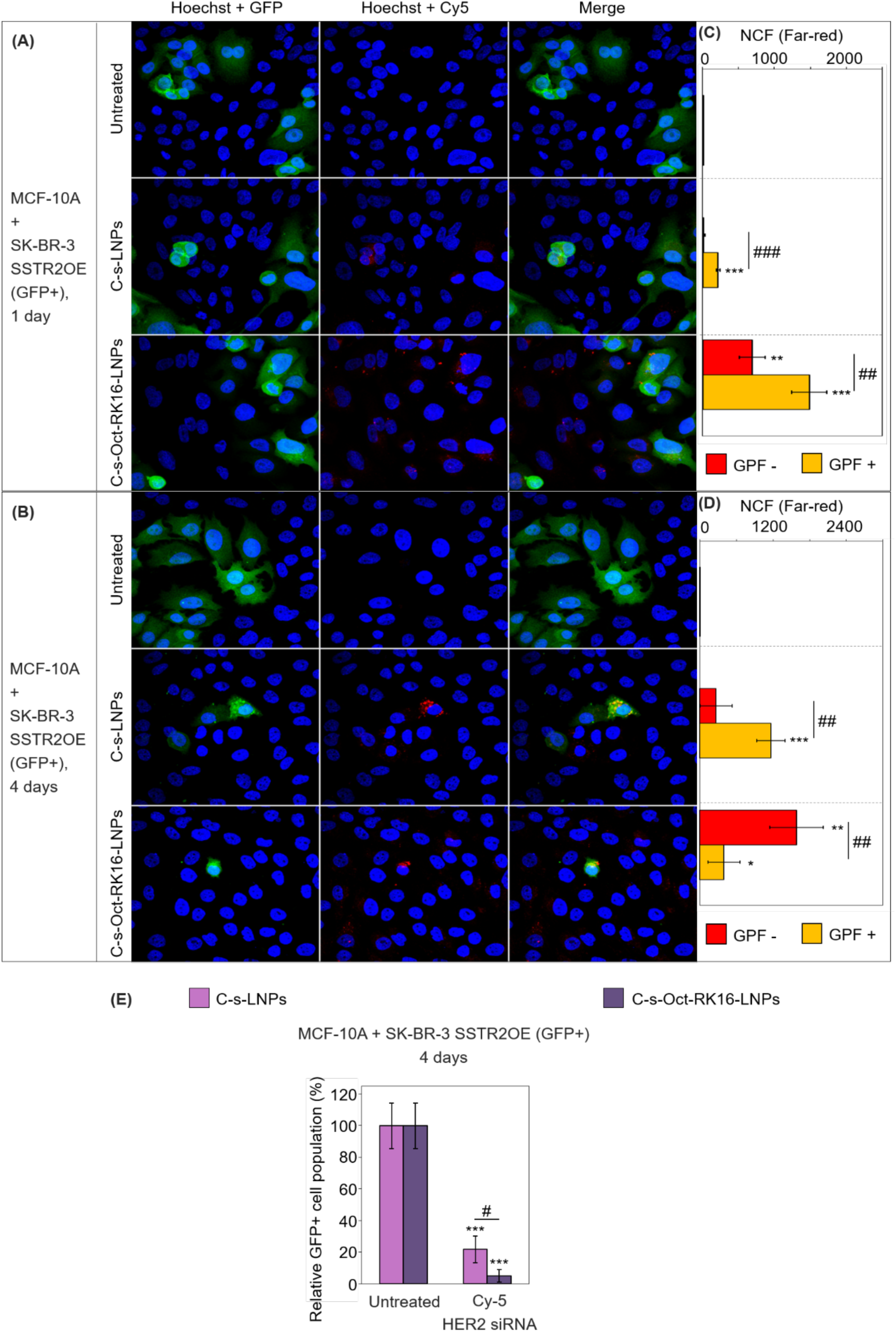
(A, B) Confocal microscopy images of co-cultures consisting of SK-BR-3 SSTR2OE (GFP+) and non-tumor MCF-10A cells following 24-hour (A) and 96-hour (B) incubation with either naked or dually-functionalized LNPs loaded with Cy5-labeled HER2 siRNA (C-s-LNPs and C-s-Oct-RK16-LNPs, respectively). Untreated cells served as negative controls. (C, D) Quantification of Cy5 fluorescence intensity in GFP-cells (MCF-10A, red) and GFP+ cells (SK-BR-3 SSTR2OE, yellow, indicating co-localization of Cy5 and GFP signals) based on images shown in panels (A) and (B). (E) Quantification of GFP fluorescence intensity normalized to blue (Hoechst) fluorescence, corresponding to the selectivity assays shown in panel (D). In all experiments, the total siRNA concentration was 20 nM. Results were normalized to untreated controls. Independent experiments were performed and quantified (n = 3). Data are expressed as mean ± standard deviation. Unpaired Student’s *t*-tests were used to compare each sample to the untreated control and between selected samples. Symbols: * / # p < 0.05; ** / ## p < 0.01; *** / ### p < 0.001. Asterisks (*) denote significance versus untreated control; hash symbols (#) indicate significance versus experimental groups.

However, at 96 hours post-treatment with dually-decorated C-s-Oct-RK16-LNPs, an apparent accumulation of Cy5-labeled siRNA was observed within MCF-10A cells (Fig. 7 B, D and Fig. S10 B, D) –an effect not observed with the naked C-s-LNP formulation. This phenomenon is likely due to the selective eradication of cancer cells by the targeted delivery of HER2 siRNA, resulting in a relative enrichment of Cy5-labeled siRNA in the remaining non-tumor population.

Consistently, long-term treatment with both LNP formulations –C-s-Oct-RK16-LNPs and naked C-s-LNPs– led to a significant reduction in the proportion of GFP+ cancer cells. Notably, the dually-functionalized C-s-Oct-RK16-LNPs induced a more pronounced cytotoxic effect. Quantification of GFP fluorescence intensity, normalized to blue (Hoechst) fluorescence and corresponding to the selectivity assays shown in Fig. 7D and Fig. S10D, revealed that C-s-Oct-RK16-LNPs achieved a (94 ± 4) % and (94 ± 3) % reduction in GFP+ cell abundance in MCF-10A + SKBR3 SSTR2OE (GFP+) (Fig. 7E) and MCF-10A + BT-474 SSTR2OE (GFP+) (Fig. S10E) co-cultures, respectively. In contrast, the naked c-s-LNPs produced a significantly lower decrease in GFP+ cell abundance [(78 ± 9) % and (68 ± 4) %, in MCF-10A + SKBR3 SSTR2OE (GFP+) (Fig. 7E) and MCF-10A + BT-474 SSTR2OE (Fig. S10E) co-cultures, respectively].

## 3. CONCLUSION

We present here a novel class of LNP formulation that might become a versatile and innovative delivery platform for ON-based targeted therapies. In contrast to currently approved LNP systems, which lack tumor selectivity, this dual functionalization approach overcomes a key limitation in ON delivery. The inclusion of both targeting (Oct) and cell penetrating (RK16)-PEG-lipid components –synthesized via modular click chemistry using azido-peptide and DBCO-PEG-lipid precursors– offers design flexibility and facilitates the incorporation of diverse targeting peptides. This adaptability supports customization for specific tumor types and organs, broadening the applicability of our platform beyond breast cancer to other malignancies and even non-cancerous diseases localized to specific tissues.

Furthermore, the co-delivery of Hsp27 and HER2 siRNAs validates the platform’s capability for combinatorial ON therapy. This enables potential applications in delivering diverse combinations of siRNAs or ASOs tailored to different oncogenic pathways or resistance mechanisms.

Future variants of this platform may incorporate alternative siRNA cargoes targeting resistance pathways in other tumor types, as well as peptides designed to engage different tumor-specific receptors. Such adaptability underscores the broad translational potential of our system to improve targeted delivery and overcome therapeutic resistance across a spectrum of malignancies.

Finally, the increased sensitivity observed in palmitate-responsive cell lines such as SK-BR-3 cells may offer an added therapeutic benefit, further enhancing the efficacy of this delivery strategy for particular tumor subtypes.

## 4. METHODS

### 4.1. Lipids, nucleic acids, and peptides

DLin-MC3-DMA [(6Z,9Z,28Z,31Z)-heptatriaconta-6,9,28,31-tetraen-19-yl-4-(dimethylamino)-butanoate] was purchased from Tebubio; cholesterol from Merck; DPG-PEG (1,2-dipalmitoyl-*rac*-glycero-3-methylpolyoxyethylene-2000) from NOF America Corporation; DSPC (distearoylphosphatidylcholine) and DBCO-DSPE-PEG {1,2-distearoyl-*sn*-glycero-3-phosphoethanolamine-*N*-[dibenzocyclooctyl(polyethyleneglycol)-2000]} from Avanti Lipids. siRNAs targeting HER2 (5’ AAG CCU CAC AGA GAU CUU GdTdT 3’) and Hsp27 (5’ UGA GAC UGC CGC CAA GUA AdTdT 3’), as well as a control scrambled siRNA (5′ AUC AAA CUG UUG UCA GCG CUG dTdT 3′), were obtained from Integrated DNA Technologies. 5’-FAM- and 5’-Cy5-labeled siRNAs targeting HER2 and Hsp27 (HPLC purified) were sourced from Metabion. The azido-derivatives of the receptor-targeting peptide octreotide (N_3_-OCT: N_3_-FCFWKTCT) and the cell-penetrating peptide penetratin (N_3_-RK16: N_3_-RQIKIWFQNRRMKWKK) were purchased from GenScript.

### 4.2. Lipid nanoparticle preparation

DSPC, DLin-MC3-DMA, cholesterol and DPG-PEG were dissolved in ethanol to prepare 10 mg/mL stock solutions. siRNAs were prepared by incubating equimolar amounts of guide and sense strands in annealing buffer (100 mM potassium acetate, 30 mM HEPES-KOH pH 7.4, 2 mM magnesium chloride) at 95 ⁰C for 5 min, followed by 1 h at 37 ⁰C. To formulate siRNA-loaded lipid nanoparticles (LNPs), the annealing buffer in the siRNA solution was exchanged with 20 mM citrate buffer (pH 4) using an Amicon centrifugal filter (Merck Millipore, 3 kDa molecular weight cut-off). A lipid mixture (10:50:38.5:1.5 molar ratio of DSPC:DLin-MC3-DMA:cholesterol:DPG-PEG) was prepared in ethanol. The lipid mixture was added dropwise to the siRNA solution after 5 min incubation at 65 ⁰C. The resulting mixture (3:2 [vol/vol] citrate buffer:ethanol, 1:16.7 [wt/wt] siRNA:lipids) was incubated at 65 ⁰C for 1 h under vigorous stirring (1600 rpm). The preparation was then sequentially extruded through 400 nm and 100 nm pore-size polycarbonate membranes (minimum 31 passes each) using an extruder (Avanti Lipids) connected to a heating block at 65 ⁰C. Finally, the LNPs were dialyzed against 20 mM citrate buffer (pH 4) for 2 h (7 kDa molecular weight cut-off) to remove ethanol, followed by 20 mM HBS buffer (pH 7.4) overnight (7 kDa molecular weight cut-off) to neutralize the ionizable lipid.

### 4.3. Lipid-peptide conjugates synthesis

In two separate reactions, 0.5 mg of each azido-peptide (N_3_-OCT and N_3_-RK16) were dissolved in 2mM DBCO-DSPE-PEG in 70% ethanol, achieving a final concentration of 2 mM for all reagents. The reaction mixtures were vortexed thoroughly and stirred overnight (400 rpm) to form the conjugates Oct-DSPE-PEG (where Oct is the receptor-targeting peptide) and RK16-DSPE-PEG (where RK16 is the cell-penetrating peptide). The crude products were directly used to functionalize the LNPs without further purification.

### 4.4. Lipid-peptide conjugates characterization

The molar absorptivities of the lipid and peptides at 309 nm were determined by measuring the absorbance at varying concentrations in 70% ethanol. The conjugation yield was calculated by monitoring the absorbance decrease at 309 nm as DBCO reacts with the azido group of the peptide. Conjugate formation was analyzed by polyacrylamide gel electrophoresis (PAGE). One nmol of DBCO-containing lipid, azido-peptides, or reaction mixtures were diluted in distilled H_2_O with 5X SDS-containing loading buffer (final volume of 10 µL), heated at 95 ⁰C for 5 min, then loaded onto a 20% gel and run in tris-glycine buffer at 100 V for 3 h. Bands were visualized by overnight staining with a Coomassie Brilliant blue G-250 solution (Bio-Rad) and imaged using a ChemiDoc Touch Imaging System (Bio-Rad). Formation of the conjugates was further confirmed by gel permeation chromatography (GPC) using an HPLC system with a PDA detector set to 220 nm (210-350 nm range) and a Ultrahydrogel 250 column (Waters) with molecular weight range of 1-80 kDa. DBCO-containing lipid, azido-peptides, and conjugation reaction mixtures were diluted in 70% ethanol to a final concentration of 100 mg/mL, and standards (1350 Da, 6500 Da, and 12000 Da) were diluted similarly. 50 µL aliquots of each standard or sample were injected into the column, with a 60:40 [vol/vol] 0.1% TFA in water:CH_3_CN mixture as the eluent (500 µL/min flow rate, 16-min runtime). ESI-TOF spectra of the conjugates were acquired using a LC/MSD TOF mass spectrometer (Agilent Technologies) with a 175 V fragmentor. Samples (1 nmol aliquots of the reaction mixtures) were diluted 1:30 in methanol and eluted with a mixture of 1:1 [vol/vol] water:CH_3_CN mixture (200 µL/min flow rate). MALDI-TOF spectra were acquired using a 4800 MALDI-TOF/TOF mass spectrometer (ABSciex) equipped with a Nd:YAG laser (355 nm wavelength, 3-7 ns pulse width, 200 Hz firing rate). Samples were diluted in a matrix containing 2,4,6-trihydroxyacetophenone (50 mg/mL in 1:1 [vol/vol] water:CH_3_CN) and ammonium citrate (50 mg/mL in water).

### 4.5. Lipid nanoparticle functionalization

Functionalized lipid nanoparticles were prepared using the post-insertion method. Oct-DSPE-PEG and/or RK16-DSPE-PEG, representing 25% [mol] of the total PEG-lipid in the naked nanoparticle formulation, were transferred to a vial, and the solvent was evaporated using a SpeedVac vacuum concentrator. The corresponding volume of preformed naked LNPs in 20 mM HBS buffer (pH 7.4) was added to the vial with the dried lipid-peptide conjugates. The mixture was thoroughly mixed and incubated at 45 ⁰C for 1 h under stirring (400 rpm) to obtain the decorated LNPs via micellar transfer.

### 4.6. Lipid nanoparticle characterization

The average size and polydispersity index (PDI) of LNPs and FLNPs were measured by dynamic light scattering using a Zetasizer Nano (Malvern Instruments) with Zetasizer 8.01 software. A 1:10 dilution of the LNPs or FLNPs in 20 mM HBS (pH 7.4) was analyzed in a low-volume quartz cuvette, with the following settings: material refractive index 1.4, absorption 0.001, dispersant refractive index 1.33, dispersant viscosity 0.8872 cP, and a 173⁰ measurement angle. All samples were analyzed in triplicate with 12-15 runs per replicate.

The surface charge (zeta potential) was measured similarly using a 1:100 dilution of the LNPs or FLNPs in 20 mM HBS (pH 7.4) in a folded capillary cell. The settings included: material refractive index 1.4, absorption 0.001, dispersant refractive index 1.33, dispersant viscosity 0.8872 cP, and dielectric constant 75.5. Triplicate measurements were made, with up to 100 runs per replicate. Additionally, fractions of both naked and decorated nanoparticle formulations were stored at 4 ⁰C, and the size and average size, PDI and zeta-potential were measured on days 3, 5, and 7 to evaluate the effects of the storage conditions on physicochemical properties and colloidal stability.

### 4.7. Encapsulation efficiency

RNA encapsulation efficiency was determined using a Qubit 4 fluorimeter and a microRNA assay kit (Thermo Fisher Scientific) for short RNAs quantitation. RNA concentrations were calculated from a standard curve generated using samples of known concentration, with the assay reagent binding to RNA and producing a fluorescent signal proportional to its concentration.

For each nanoparticle formulation, a 1:5 dilution in 20 mM HBS buffer (pH 7.4) was added to the assay reagent to determine unencapsulated RNA concentration, and a 1:5 dilution in 0.1% Triton X-100 in 20 mM HBS (pH 7.4) was used to determine total RNA concentration. Encapsulated RNA concentration was calculated by subtracting the unencapsulated RNA concentration from the total RNA concentration. Encapsulation efficiency was expressed as the ratio of encapsulated RNA to total RNA, as a percentage.

Additionally, fractions of the both naked and decorated nanoparticle formulations were stored at 4 ⁰C, and encapsulation efficiency was re-assessed on days 3, 5 and 7 to evaluate the effects of the storage conditions on physicochemical properties and colloidal stability.

### 4.8. Cell culture

SKBR3, BT-474, HEK 293T and MCF10A cells were obtained from ATCC, while SKBR3 SSTR2OE (GFP+) and BT-474 SSTR2OE (GFP+) cells were generated in-house. SKBR3 and SKBR3 SSTR2OE (GFP+) cells were cultured in McCoy’s 5A medium (Gibco) supplemented with 10% fetal bovine serum, 100 U/mL penicillin and 100 µg/mL streptomycin. BT-474 and BT-474 SSTR2OE (GFP+) cells were maintained in Dulbecco’s Modified Eagle Medium F12 (Gibco) supplemented with 10% fetal bovine serum, 100 U/mL penicillin and 100 µg/mL streptomycin. HEK 293T cells were cultured in Dulbecco’s Modified Eagle Medium (Gibco) supplemented with 10% fetal bovine serum, 100 U/mL penicillin and 100 µg/mL streptomycin. MCF10A cells were maintained in Dulbecco’s Modified Eagle Medium F12 (Gibco) supplemented with 10% fetal bovine serum, 100 U/mL penicillin, 100 µg/mL streptomycin, 10 µg/mL human insulin, 12.5 ng/µL epidermal growth factor, 250 ng/mL hydrocortisone and 100 ng/mL cholera toxin. All cell lines were cultured at 37°C in a humidified atmosphere with 5% CO_2_.

### 4.9. Generation of SSTR2-overexpressing HER2+ breast cancer cell lines

SKBR3 and BT-474 cells stably overexpressing somatostatin receptor type 2 (SSTR2) and green fluorescent protein (GFP) were generated by viral transduction. HEK 293T cells were seeded in 150 mm dishes at a density of 4 million cells per dish in medium without antibiotics and co-transfected with three plasmids for lentiviral particle production: psPAX2 (packaging plasmid; Addgene), pMD2.G (VSV-G envelope-expressing plasmid; Addgene), and pGenLenti (transfer vector containing an insert of the SSTR2 and the GFP genes separated by the ribosomal-skipping inducing peptide P2A; GenScript). Cells were co-transfected with 10 µg of psPAX2, 5 µg of pMD2.G and 15 µg of pGenLenti per dish, with 1 mg/mL polyethylenimine (1:1 [wt/wt] DNA:PEI). After 5 hours, the medium was replaced with fresh complete medium. Viral particles were harvested 48 and 72 hours after transfection by filtering the culture media through a 0.45 µm PES filter, and polybrene was added to a final concentration of 1 µg/mL to enhance infection efficiency.

SKBR3 and BT-474 cells were seeded in 150 mm dishes at a density of 3 million cells per dish in complete media, and an additional plate per cell line was prepared as a non-infected negative control for antibiotic selection. Cells were infected with filtered viral harvests 24 and 48 hours after seeding. Six hours after the second infection, media were replaced with fresh complete media supplemented with 4 µg/mL puromycin. After 72 hours, viral transduction efficiency was assessed by fluorescence microscopy for GFP expression and cell survival, and the selection media were replaced with fresh complete media after verifying the death of the uninfected control cells.

### 4.10. Flow cytometry

SSTR2 and GFP expression in SKBR3, SKBR3 SSTR2OE (GFP+), BT-474, BT-474 SSTR2OE (GFP+) and MCF10A cells was assessed by flow cytometry. 10^6^ cells were harvested, incubated for 30 min at room temperature with an APC-labeled anti-SSTR2 monoclonal antibody (R&D Systems; 1:20 dilution in PBS) or PBS as a control, washed, and resuspended in 5 µg/mL DAPI (in PBS) for viability assessment. Cells were stored at 4 ⁰C until further analysis. Fluorescence was measured using a Gallios flow cytometer (Beckman Coulter), analyzing 15000 cells per sample, with fluorescence intensity displayed on a logarithmic scale.

To evaluate the selectivity of octreotide-decorated LNPs for SSTR2-overexpressing (GFP+) cells, 1:1 co-cultures of GFP+ and GFP-cells (75000 cells per well) were seeded on 24-well plates in complete media and incubated at 37 ⁰C in a humidified atmosphere with 5% CO_2_. After overnight culture, cells were transfected with nanoparticle formulations at various doses (total volume of 0.5 mL) or left untreated as controls. After 48 hours (SKBR3 + SKBR3 SSTR2OE co-cultures) or 72 hours (BT-474 + BT-474 SSTR2OE co-cultures), cells were harvested, resuspended in a 1:1 mixture of complete media without phenol red and 5 µg/mL DAPI in PBS (total volume of 250 µL), and transferred to flow cytometry tubes for analysis. Fluorescence was measured using an Aurora 4L cytometer (Cytek Biosciences; equipped with SpectroFlo software), analyzing 15000 events per sample. The populations of live cells (DAPI-negative) and SSTR2-overexpressing (GFP-positive) cells were quantified. Cell viability was expressed as a percentage of untreated controls, and LC50 values (median-effect doses corresponding to a 50% reduction in cell viability) were calculated using CompuSyn software (ComboSyn). All experiments were performed in triplicate.

### 4.11. Fluorescence microscopy

Cell uptake of naked and decorated LNPs was evaluated by fluorescence confocal microscopy. Cells were seeded at a density of 50000 cells per well (or 25000 cells of each line per well in co-cultures) on 24-well plates with 12 mm cover glasses at the well bottom in complete media and incubated at 37 ⁰C in a humidified atmosphere with 5% CO_2_. After overnight culture, cells were transfected with the corresponding nanoparticle formulations at different doses (0.5 mL total volume) or left untreated as a control. Following 24, 48, or 96 hours of incubation period, cells were fixed with 4% paraformaldehyde at pH 7.4 for 10 min at room temperature, permeabilized with 0.5% Triton X-100 in PBS for 5 min, and incubated in 1 µg/mL Hoechst for 5 min for nuclear staining. The cover slides were then removed, mounted on slides with Fluoromount-G (Electron Microscopy Sciences) and incubated overnight at 4 ⁰C before imaging. Fluorescence of GFP, nuclear staining and siRNAs was observed using an SPE confocal microscope (Leica) with LAS AF software. Hoechst fluorescence (461 nm emission) was visualized with a 405 nm laser (10% intensity, 1000 gain), GFP and fluorescein fluorescence (508 nm and 517 nm emission, respectively) were visualized with a 488 nm laser (15% intensity, 1000 gain), and Cy5 fluorescence (667 nm emission) was visualized with a 635 nm laser 15% intensity, 1000 gain). The images were analyzed using ImageJ software (NIH) and the corrected total cell fluorescence was calculated for each sample and fluorophore, and the values were normalized relative to the untreated control.

### 4.12. Viability assay

The effect of naked and decorated LNPs on cell growth rate was assessed using the crystal violet assay. Cells (75000 per well) were seeded in 24-well plates in complete media. After overnight incubation, the cells were transfected with the corresponding nanoparticle formulations at varying doses (total volume of 0.5 mL) or left untreated (negative control). Following a 72-hour or 96-hour incubation period, cells were fixed with 4% paraformaldehyde (pH 7.4) for 15 min at room temperature. They were then stained with freshly prepared 4.5% crystal violet for 20 min at room temperature, thoroughly washed with distilled water, and incubated with 10% acetic acid for 20 min at room temperature under gentle rocking to dissolve the dye. The resulting samples were diluted fourfold with distilled water and transferred to 96-well plates. Absorbance was measured at 570 nm using a BioTek ELx808 microplate reader (Agilent Technologies) with Gen5 2.09 software. Cell viability was expressed as a percentage relative to the untreated controls, and the corresponding LC50 values and combination indices (Cis) [quantitative measures that define drug interactions as synergistic (CI < 1), antagonistic (CI > 1) or additive (CI = 1)] were calculated using CompuSyn software (ComboSyn). All experiments were performed in triplicate.

### 4.13. Cell lysis and western blot

ErbB2 and Hsp27 protein knockdown by naked and decorated LNPs was analyzed by western blot. Cells (100000 per well) were seeded on 24-well plates in complete media. After overnight incubation, the cells were transfected with the corresponding nanoparticle formulations at varying doses (total volume of 0.5 mL) or left untreated (negative control). Following a 72 h incubation period, cells were lysed by scraping on ice in RIPA lysis buffer with protease inhibitors (Roche) and incubated at 4 ⁰C for 30 min under stirring. Lysates were then clarified by centrifugation at 13000 rpm for 30 min at 4 ⁰C and protein concentration was determined via DC Protein Assay (Bio-Rad). Next, 30 µg of protein per sample were resolved by SDS-PAGE (tris-glycine running buffer, 140 V, 1 h) and transferred onto a polyvinylidene difluoride membrane (Bio-Rad). The membrane was blocked with 5% skim milk in TBS with 0.1% Tween-20 for 1 h at room temperature under rocking and then probed with polyclonal rabbit antibodies against ErbB2 (Cell Signaling Technologies, 1:1000 dilution in 5% BSA in TBS 0.1% Tween-20) and Hsp27 (Abcam, 1:1000 dilution in 5% BSA in TBS 0.1% Tween-20) overnight at 4 ⁰C. The membrane was then probed with an anti-rabbit (goat) IgG HRP-conjugated antibody (Thermo Fisher Scientific, 1:5000 dilution in 5% BSA in TBS 0.1% Tween-20) for 1 h at room temperature and with an anti β-actin HRP-conjugated antibody (Abcam, 1:20000 dilution in 5% BSA in TBS 0.1% Tween-20) for 1 h at room temperature. The membrane was revealed with chemiluminescent Clarity Western ECL Blotting Substrate (Bio Rad) using a ChemiDoc Touch Imaging System (Bio-Rad). The images were analyzed with ImageJ software (NIH software) and protein levels were expressed as a percentage of the controls. All experiments were performed in triplicate.

### 4.14. Statistical analysis

Where appropriate, and unless otherwise stated, results are indicated as mean ± standard deviation (SD). Statistical differences were determined using two-tailed Student’s t-tests for unpaired observations, considering p-values lower than 0.05 to indicate significance.

## ACKNOWLEDGEMENTS

This work was supported by the Instituto de Salud Carlos III (PI18/01964 to M.T.), by the Spanish Ministry of Science, Innovation and Universities through the National Program FPU (grant number FPU19/05307 to P.M.), the Spanish Ministry of Science (PDI2021-122478NB-I00, PCI2022-134976-2 and PLEC2024-011123),, the Instituto de Salud Carlos III (XNA-Hub Project) and the Generalitat de Catalunya (AGAUR. Ref.: 2021SGR0086) all granted to M.O’s group. We thank the Mass Spectrometry Unit of the Faculty of Chemistry, the Flow Cytometry Unit of the Barcelona Science Park and the Separative Techniques Unit of the Barcelona Science Park (University of Barcelona; CCiTUB) for data acquisition and technical support. We also thank the Advanced Digital Microscopy Core Facility of the IRB Barcelona for help and valuable comments and the advice of Dr. I. Brun-Heath at the EBL.

