## Supplementary Information for "Dually-decorated palmitate-containing lipid nanoparticles for the targeted delivery of siRNAs against HER2 and Hsp27 in HER2+ breast cancer"

**Electronic Supplementary Information**

### Table of contents

Figure S1 ........ S3
Figure S2 ........ S4
Figure S3 ........ S5
Figure S4 ........ S6
Figure S5 ........ S7
Figure S6 ........ S8
Figure S7 ........ S9
Figure S8 ........ S10
Figure S9 ........ S11
Figure S10 ........ S12
Figure S11 ........ S14

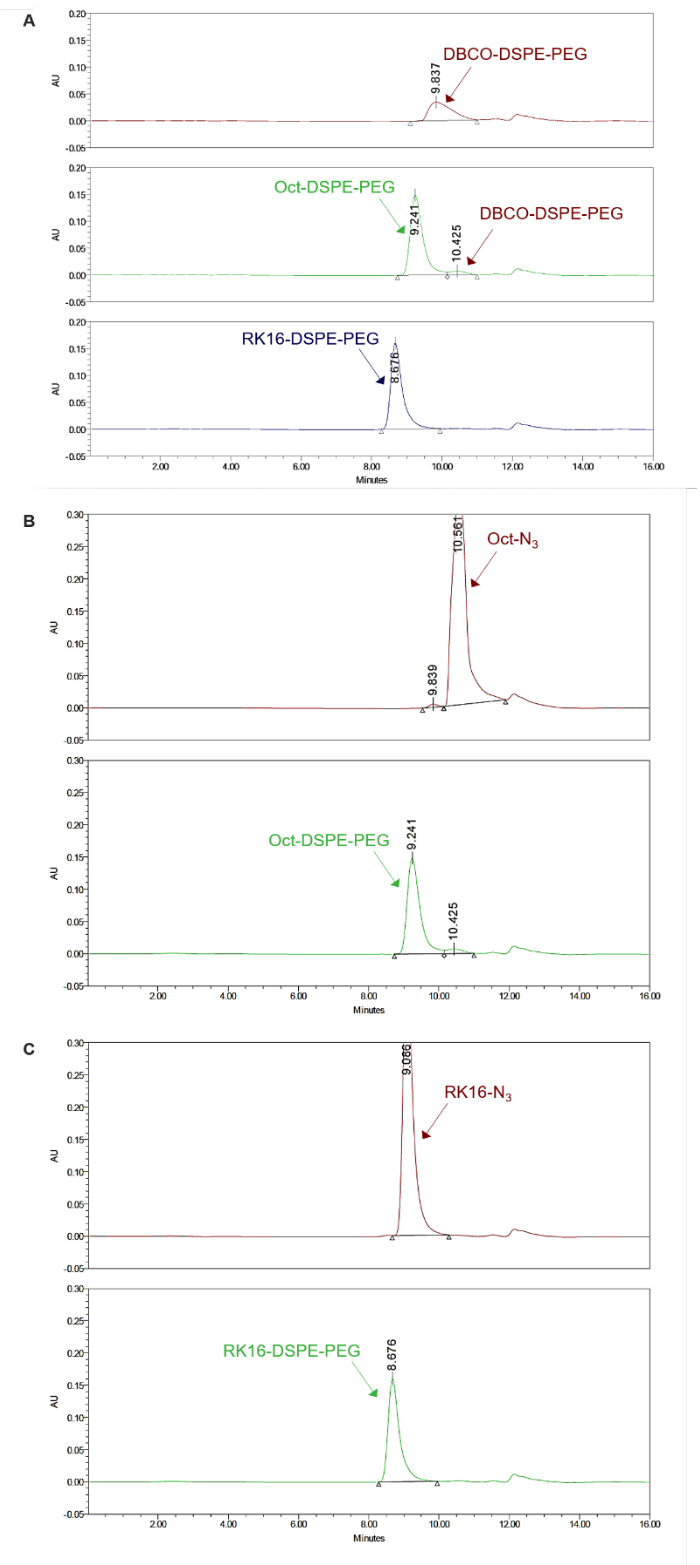


**Figure S1.** Gel permeation chromatography (GPC) analysis of the reaction between DBCO-DSPE-PEG and RK16-$N_3$ or Oct-$N_3$. (A) Comparison of the mobilities of the conjugates RK16-DSPE-PEG and Oct-DSPE-PEG with the starting cyclooctyne DBCO-DSPE-PEG. (B) GPC profile of Oct-DSPE-PEG and the starting azide Oct-$N_3$. (C) GPC profile of RK16-DSPE-PEG and the starting azide RK16-$N_3$.

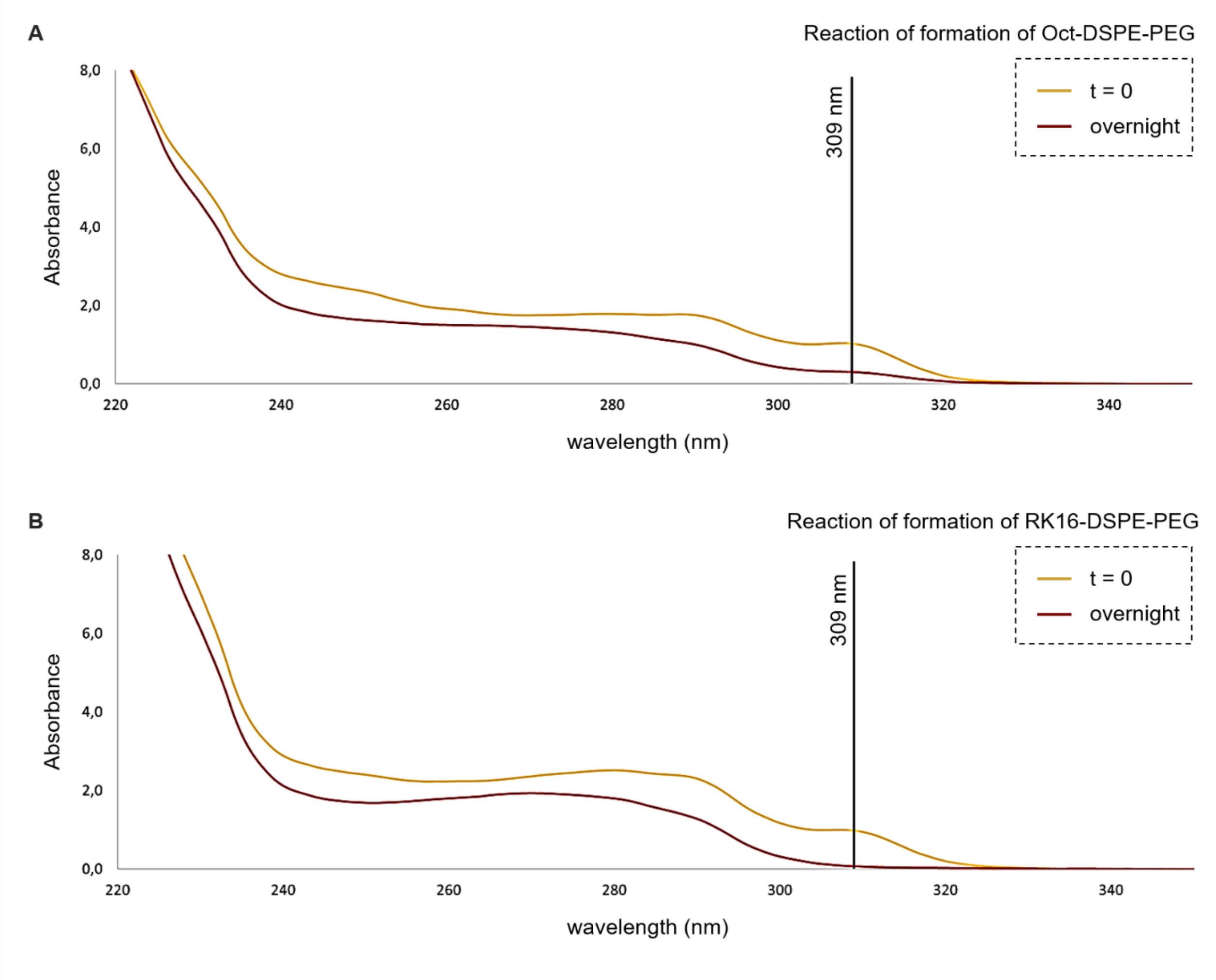


**Figure S2.** Analysis of the reaction of formation of Oct-DSPE-PEG and RK16-DSPE-PEG by UV-vis spectroscopy. Measurement of the A309 associated to the unreacted DBCO-DSPE-PEG lipid enables to indirectly quantify the resulting conjugates.

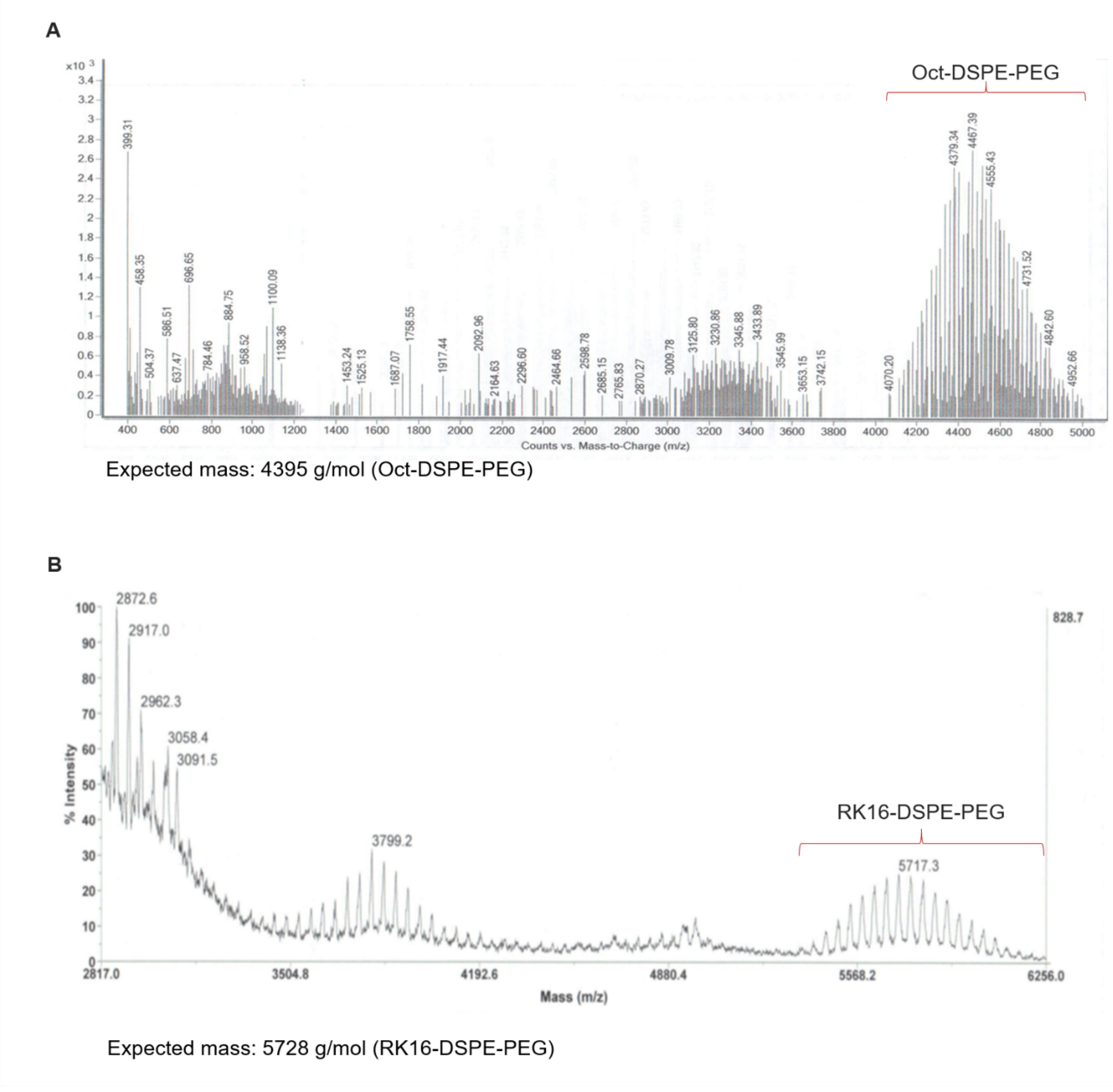


**Figure S3. (A)** ESI-TOF mass spectrometry analysis of the crude PEGylated peptide-lipid conjugate Oct-DSPE-PEG and (B) MALDI-TOF mass spectrometry analysis of the crude PEGylated peptide-lipid conjugate RK16-DSPE-PEG.

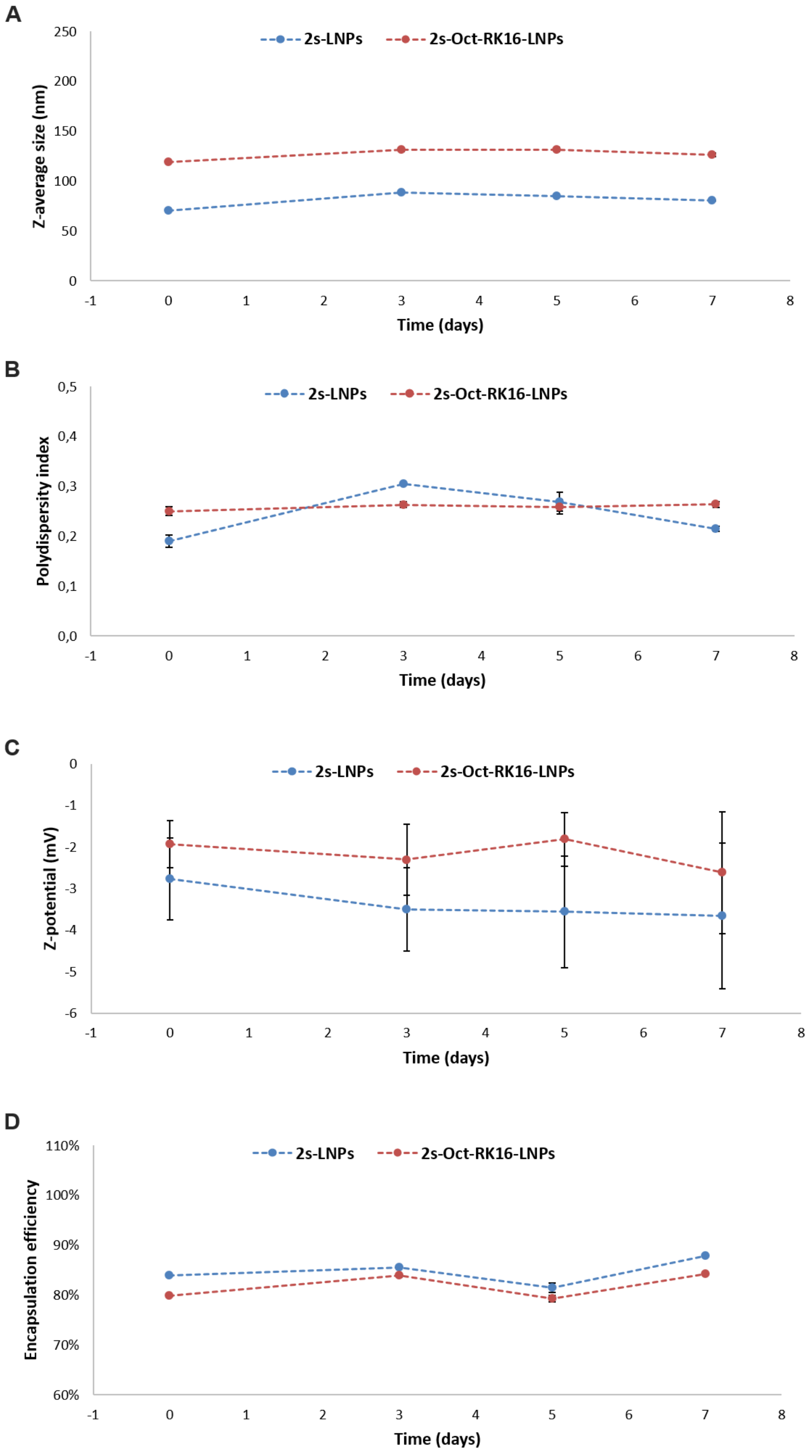


**Figure S4.** Colloidal stability of naked and dually-decorated lipid nanoparticles loaded with a 1:1 mixture of Hsp27 and HER2 siRNAs (2s-LNPs and 2s-Oct-RK16-LNPs, respectively) over a 7-day period at 4 ºC. (A) Average size, (B) polydispersity index, (C) zeta potential, and (D) encapsulation efficiency. Size measurements were taken at the indicated time points to monitor changes in size distribution, surface charge and RNA encapsulation over time.

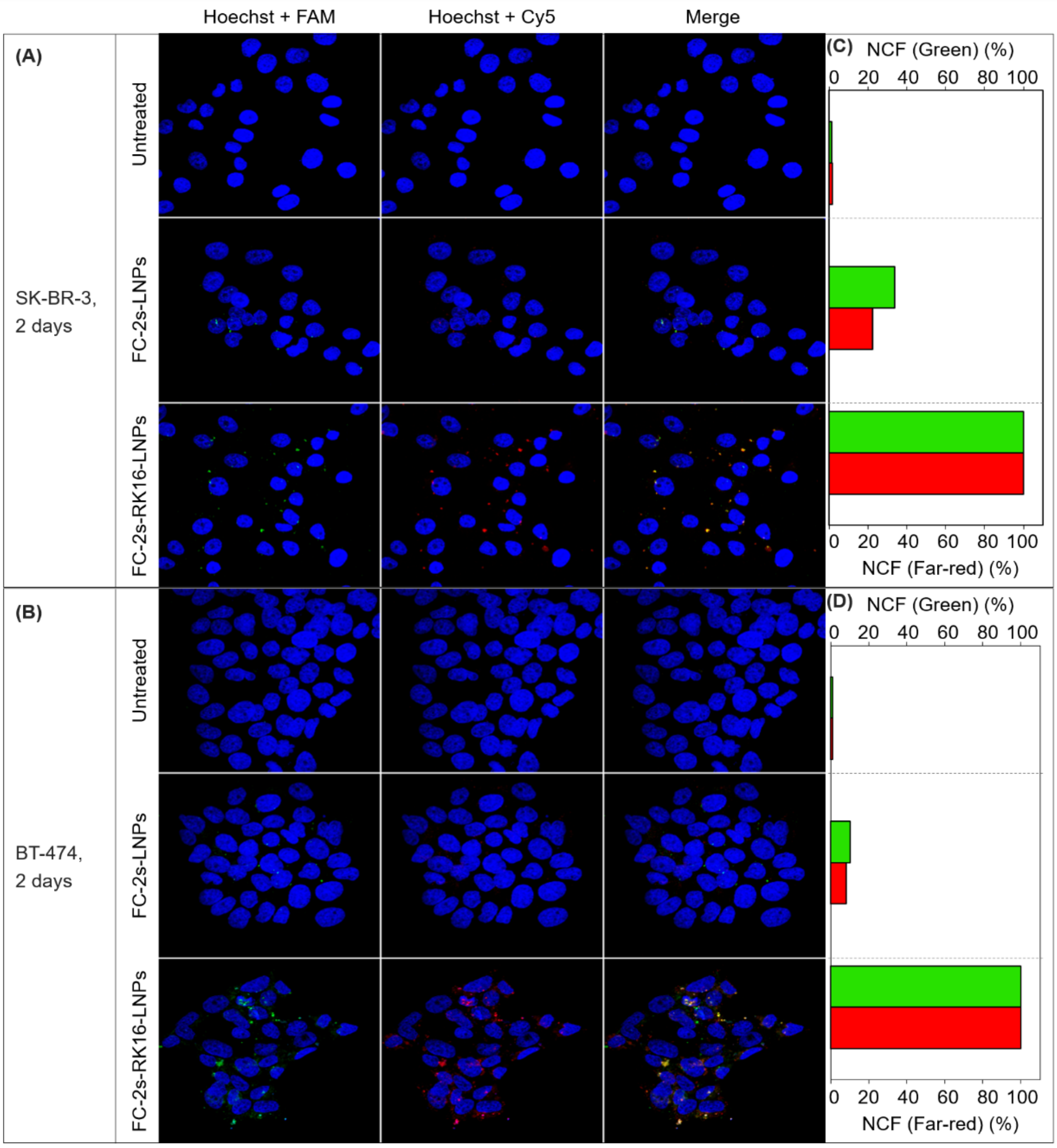


**Figure S5.** (A, B) Confocal microscopy images of SK-BR-3 (A) and BT-474 (B) cells incubated with either naked or RK16-functionalized LNPs, both loaded with a 1:1 mixture of FAM-Hsp27 siRNA and Cy5-HER2 siRNA (FC-2sLNP and FC-2s-RK16-LNPs, respectively) at a total siRNA concentration of 20 nM for 2 days. Untreated cells were used as negative controls in both cases. (C, D) Quantification of green (FAM) and red (Cy5) fluorescence intensities corresponding to the confocal images in panels (A) and (B), respectively.

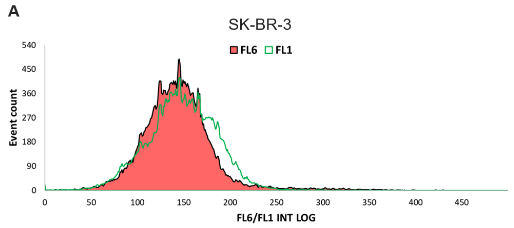


| Region | Number | % Gated | X-Mean |
|---|---|---|---|
| All | 15000 | 100.00% | 1.14 |
| **Positive** | **1966** | **13.11%** | **2.28** |

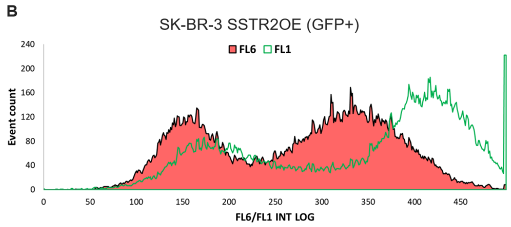


| Region | Number | % Gated | X-Mean |
|---|---|---|---|
| All | 15000 | 100.00% | 43.60 |
| **Positive** | **11789** | **78.59%** | **55.20** |

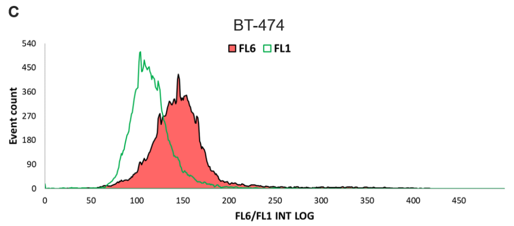


| Region | Number | % Gated | X-Mean |
|---|---|---|---|
| All | 15000 | 100.00% | 1.44 |
| **Positive** | **4477** | **29.85%** | **2.17** |

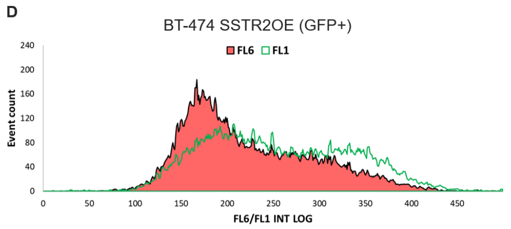


| Region | Number | % Gated | X-Mean |
|---|---|---|---|
| All | 15000 | 100.00% | 14.50 |
| **Positive** | **13001** | **86.74%** | **16.40** |

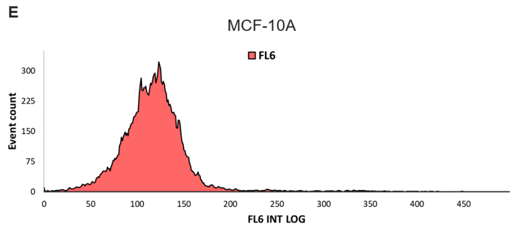


| Region | Number | % Gated | X-Mean |
|---|---|---|---|
| All | 15946 | 100.00% | 1.00 |
| **Positive** | **2498** | **15.67%** | **2.13** |

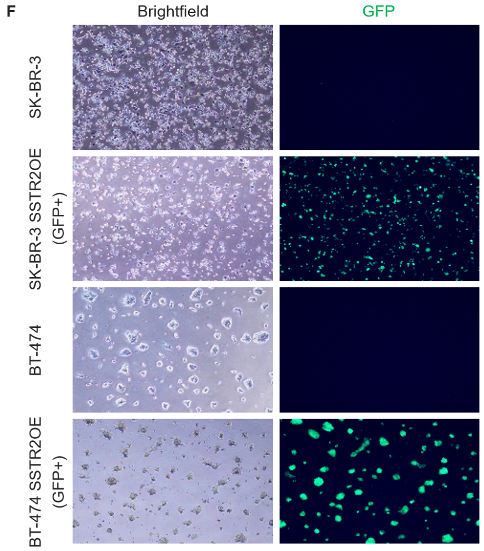


**Figure S6.** Flow cytometry analysis of SSTR2 and GFP expression in parental SK-BR-3 (A) and BT-474 (C) cells, SSTR2-overexpressing SK-BR-3 SSTR2OE (GFP+) (B) and BT-474 SSTR2OE (GFP+) (D) cells, and non-cancerous MCF-10A cells (E, analyzed for SSTR2 expression only). (F) Fluorescence microscopy images showing GFP expression in SK-BR-3, SK-BR-3 SSTR2OE (GFP+), BT-474, and BT-474 SSTR2OE (GFP+) cells. FL1: GFP fluorescence (channel 1); FL6: APC fluorescence for SSTR2 detection (channel 6); INT LOG: fluorescence intensity displayed on a logarithmic scale.

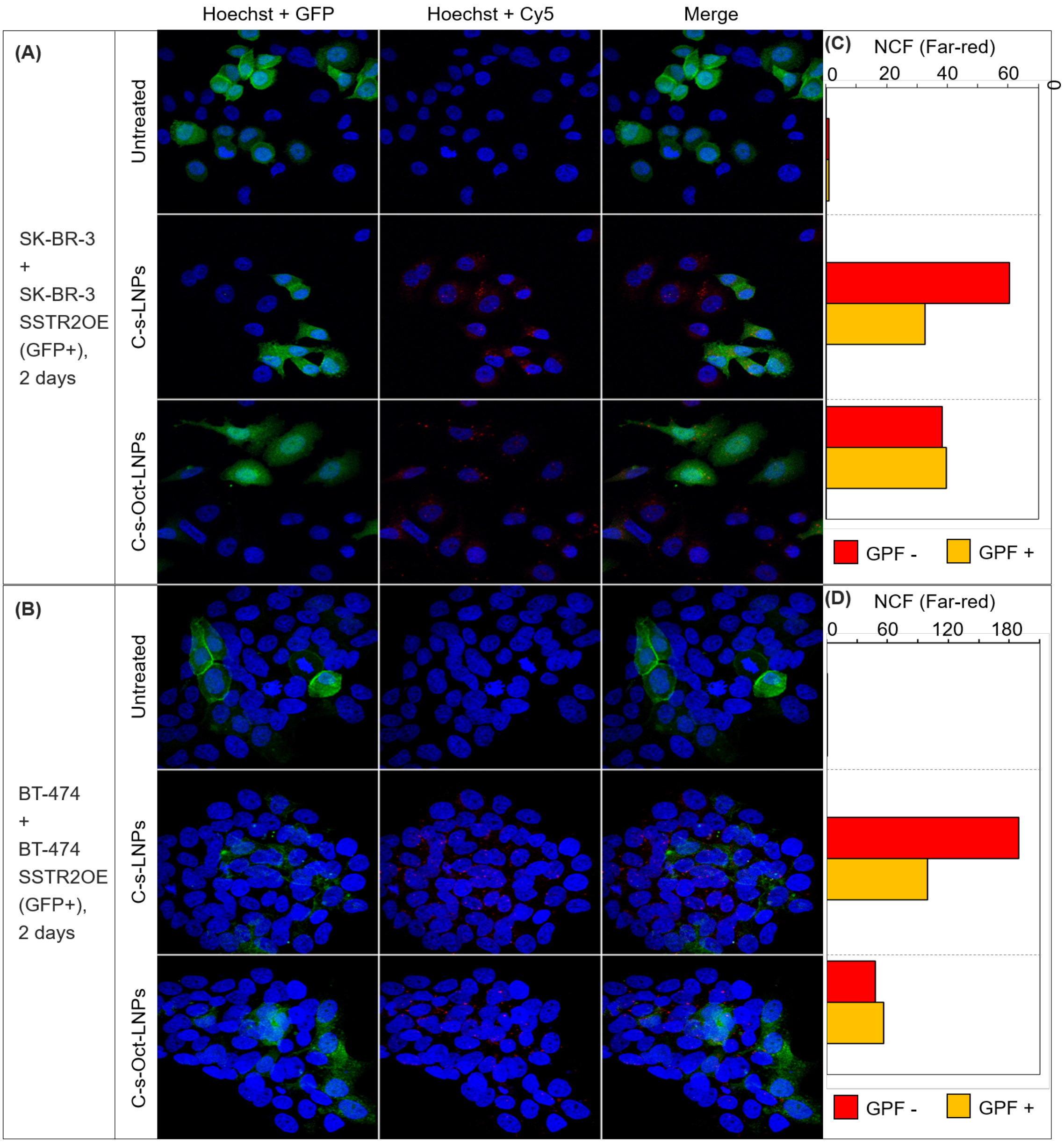


**Figure S7.** (A, B) Confocal microscopy images of co-cultures consisting of SK-BR-3 and SK-BR-3 SSTR2OE (GFP+) cells (A), and BT-474 and BT-474 SSTR2OE (GFP+) cells (B), incubated for 48 hours with either naked or Oct-functionalized LNPs, both loaded with Cy5-labeled HER2 siRNA (C-s-LNPs and C-s-Oct-LNPs, respectively) at a final siRNA concentration of 40 nM. Untreated cells served as negative controls. (C, D) Quantification of red (Cy5) fluorescence intensity in GFP- cells (red) and GFP+ cells (yellow, indicating co-localization of Cy5 and GFP signals) from images shown in panels (A) and (B).

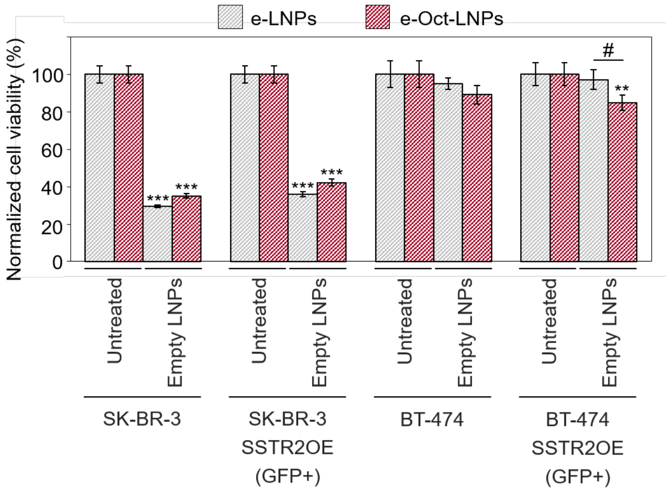


**Figure S8.** Crystal violet cell viability assay performed 96 hours post-transfection of SK-BR-3, SK-BR-3 SSTR2OE (GFP+), BT-474 and BT-474 SSTR2OE (GFP+) cells with either naked or Oct-functionalized empty LNPs (e-LNPs and e-Oct-LNPs). Results were normalized to untreated controls. All experiments were performed independently in triplicate (n = 3). Data are presented as mean ± standard deviation. Statistical comparisons were made using unpaired Student's *t*-tests against untreated control and between selected experimental groups. Symbols: # $p < 0.05$; ** $p < 0.01$; *** $p < 0.001$. Asterisks (*) indicate significance versus untreated control; hash symbols (#) indicate significance versus experimental groups.

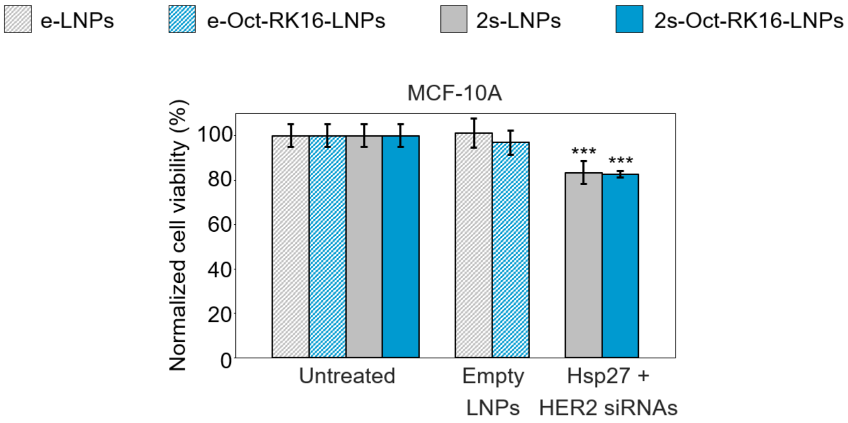


**Figure S9.** Crystal violet cell viability assay performed 96 hours post-transfection of MCF-10A cells with either naked or Oct-RK16-functionalized LNPs, formulated as either empty or loaded with a 1:1 mixture of Hsp27 and HER2 siRNAs (e-LNPs, 2s-LNPs, e-Oct-RK16-LNPs, 2s-Oct-RK16-LNPs). In all experiments, the total siRNA concentration was 20 nM. Results were normalized to untreated controls. All experiments were performed independently in triplicate (n = 3). Data are presented as mean ± standard deviation. Statistical comparisons were made using unpaired Student's *t*-tests against untreated control and between selected experimental groups. Symbols: *** $p < 0.001$. Asterisks (*) indicate significance versus untreated controls.

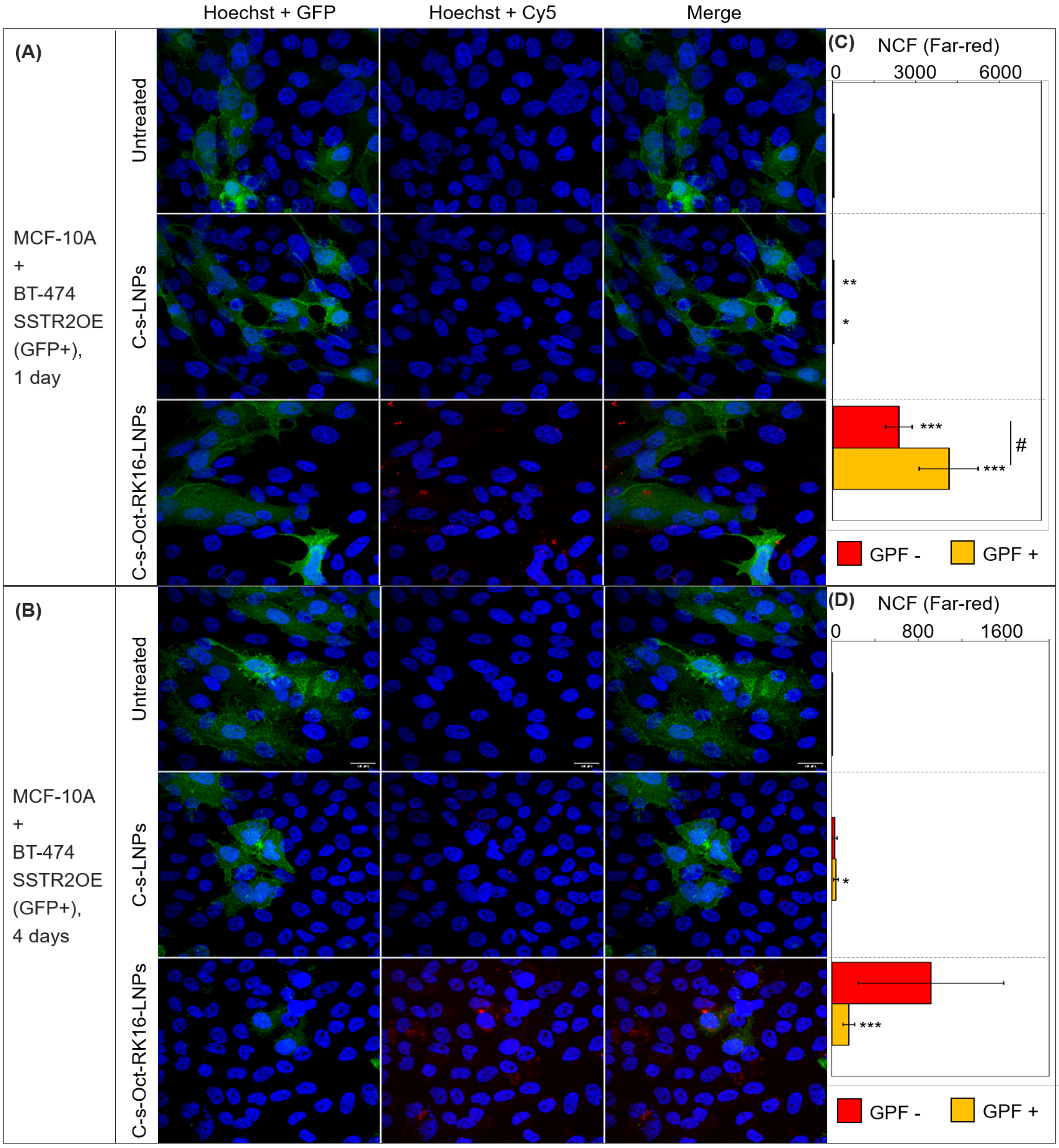
Hoechst + GFP
Hoechst + Cy5
Merge
(A)
MCF-10A
+
BT-474
SSTR2OE
(GFP+),
1 day
Untreated
C-s-LNPs
C-s-Oct-RK16-LNPs
(C)
NCF (Far-red)
0
3000
6000
**
*
***
***
#
GPF -
GPF +
(B)
MCF-10A
+
BT-474
SSTR2OE
(GFP+),
4 days
Untreated
C-s-LNPs
C-s-Oct-RK16-LNPs
(D)
NCF (Far-red)
0
800
1600
*
***
GPF -
GPF +

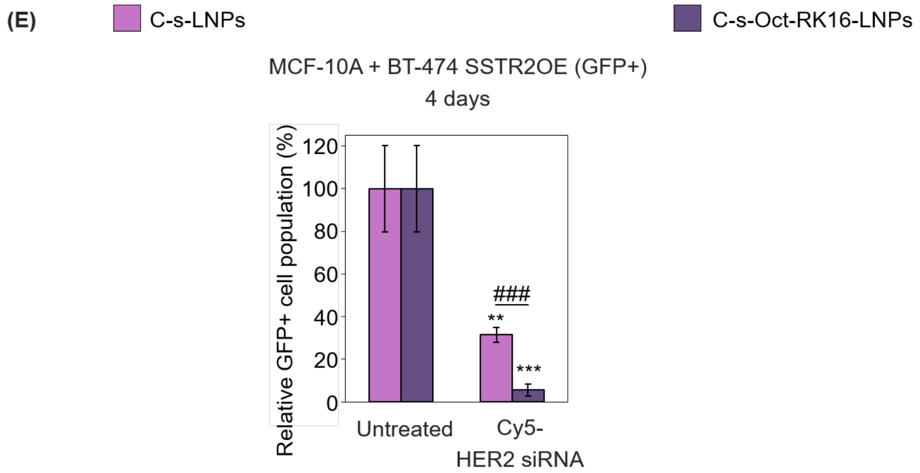
(E)
C-s-LNPs
C-s-Oct-RK16-LNPs
MCF-10A + BT-474 SSTR2OE (GFP+)
4 days
Relative GFP+ cell population (%)
120
100
80
60
40
20
0
###
**
***
Untreated
Cy5-
HER2 siRNA

**Figure S10.** (A, B) Confocal microscopy images of co-cultures consisting of BT-474 SSTR2OE (GFP+) and non-tumor MCF-10A cells following 24-hour (A) and 96-hour (B) incubation with either naked or dually-functionalized LNPs loaded with Cy5-labeled HER2 siRNA (C-s-LNPs and C-s-Oct-RK16-LNPs, respectively). Untreated cells served as negative controls. (C, D) Quantification of Cy5 fluorescence intensity in GFP- cells (MCF-10A, red) and GFP+ cells (SK-BR-3 SSTR2OE, yellow, indicating co-localization of Cy5 and GFP signals) based on images shown in panels (A) and (B). (E) Quantification of GFP fluorescence intensity normalized to blue (Hoechst) fluorescence, corresponding to the selectivity assays shown in panel (B). In all experiments [panels (A-E)], the total siRNA concentration was 20 nM. Results were normalized to untreated controls. Independent experiments were performed and quantified (n = 3). Data are expressed as mean ± standard deviation. Unpaired Student's *t*-tests were used to compare each sample to the untreated control and between selected samples. Symbols: ** $p < 0.01$; *** / ### $p < 0.001$. Asterisks (*) denote significance versus untreated control; hash symbols (#) indicate significance versus experimental groups.

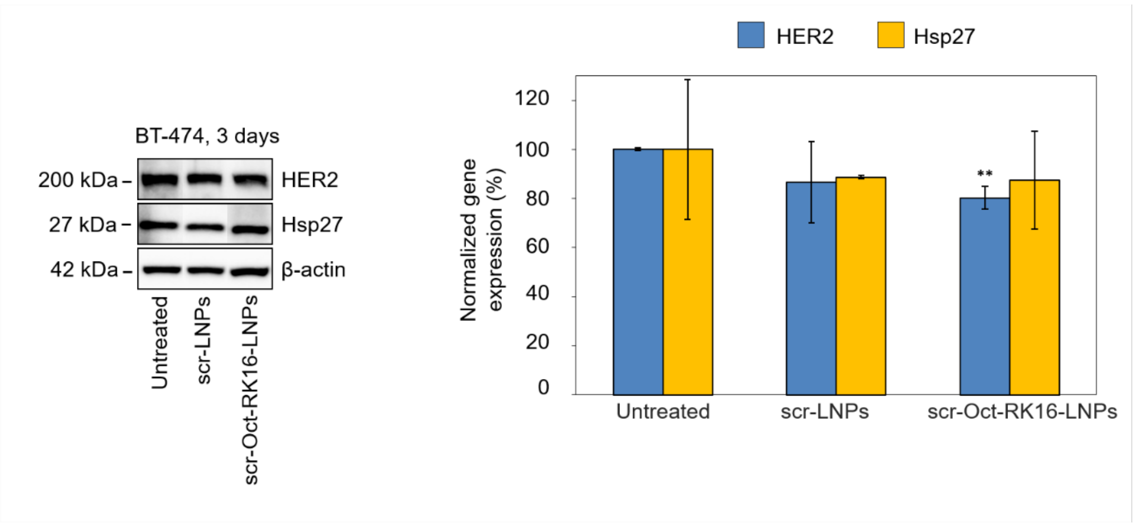


**Figure S11.** Representative immunoblot and quantitative analysis of protein expression for HER2, Hsp27 and actin (internal control) from BT-474 cells treated with scr-LNPs, and scr-Oct-RK16-LNPs for 72 hours. In all experiments, the total siRNA concentration was 20 nM. Results were normalized to untreated controls. Independent experiments were performed and quantified (n = 3). Data are expressed as mean ± standard deviation. Unpaired Student's *t*-tests were used to compare each sample to the untreated control and between selected samples. Symbols: ** $p < 0.01$. Asterisks (*) denote significance versus untreated control.
